# Lmx1a is a master regulator of the cortical hem

**DOI:** 10.1101/2022.10.25.513532

**Authors:** Igor Y. Iskusnykh, Nikolai Fattakhov, Yiran Li, Laure Bihannic, Matthew K. Kirchner, Ekaterina Y. Steshina, Paul A. Northcott, Victor V. Chizhikov

## Abstract

Development of the nervous system depends on signaling centers – specialized cellular populations that produce secreted molecules to regulate neurogenesis in the neighboring neuroepithelium. Some signaling centers also generate key types of neurons. The formation of a signaling center involves its induction, the maintenance of expression of its secreted molecules, and cell differentiation and migration events. How these distinct processes are coordinated during signaling center development remains unknown. Here we show that Lmx1a acts as a master regulator to orchestrate the formation and function of the cortical hem (CH), a critical signaling center that controls hippocampus development. Lmx1a co-regulates CH induction, its Wnt signaling, and the differentiation and migration of CH-derived Cajal-Retzius neurons. Combining RNAseq, genetic, and rescue experiments, we identified major downstream genes that mediate distinct Lmx1a-dependent processes. Our work revealed that signaling centers in the mammalian brain employ master regulatory genes and established a framework for analyzing signaling center development.

## Introduction

In the vertebrate central nervous system (CNS), tissue patterning and subsequent neurogenesis are regulated by signaling centers – localized groups of cells that produce secreted molecules that regulate development of neighboring cells (Bielen et al., 2017; Cavodeassi and Houart, 2012; Manfrin et al., 2019). Formation of a signaling center is a complex process that involves its induction, segregation of its cells from the adjacent tissue, and the initiation and maintenance of expression of secreted molecules that mediate its function. In many cases, cells that comprise signaling centers also differentiate into neurons that migrate to specific brain regions and regulate their morphogenesis and/or form neural circuits (Kiecker and Lumsden, 2012; Subramanian and Tole, 2009a). Currently, little is known about how different aspects of signaling center formation are coordinated during development. In vertebrates, development of several cell types or even organs depends on master regulatory genes – intrinsic factors that orchestrate multiple developmental processes, such as transcription factors *Atoh1* in cerebellar granule cells or *Pax6* in the eye (Baker et al., 2018; Graw, 2010; Klisch et al., 2011; Srivastava et al., 2013). It is unknown, however, whether the development of signaling centers in the CNS involves master regulatory genes.

One of the key signaling centers in the mammalian CNS is the cortical hem (CH), which develops at the telencephalic dorsal midline, between the choroid plexus and cortical neuroepithelium (Grove et al., 1998; Jones et al., 2019; Moore and Iulianella, 2021; Sindhu et al., 2019). The CH is necessary and sufficient for the development of the hippocampus in adjacent neuroepithelium (Mangale et al., 2008). Loss of CH or its signaling molecule Wnt3a leads to the near-complete loss of the hippocampus, associated with reduced proliferation in the hippocampal primordium (Lee et al., 2000). Canonical Wnt signaling from the CH (involving Wnt3a and likely other Wnts) also promotes the formation of the transhilar glial scaffold, which guides the migration of neural progenitors from the dentate neuroepithelium (DNe) in the ventricular zone to the hippocampal dentate gyrus (DG) (Zhou et al., 2004). In addition to Wnt signaling, CH gives rise to Cajal-Retzius (CR) cells, short-lived neurons that migrate to the hippocampal field, where they form the hippocampal fissure, the sulcus that separates the DG from the CA1 field. CR cells also promote the formation of the transhilar glial scaffold (Bielle et al., 2005; Causeret et al., 2021; Hodge et al., 2013; Meyer et al., 2004).

To date, intrinsic factors that confer CH fate remain elusive. For example, loss of transcription factors Dmrt3/4/5 or Gli3 compromises CH formation (De Clercq et al., 2018; Fotaki et al., 2011; Grove et al., 1998; Kikkawa and Osumi, 2021; Quinn et al., 2009; Saulnier et al., 2013). However, both *Dmrt* and *Gli3* are expressed broadly in the telencephalic medial neuroepithelium (in both the CH and beyond), and their overexpression was not reported to induce ectopic CH, suggesting that they play permissive rather than instructive roles in CH development (Subramanian et al., 2009a; Subramanian and Tole, 2009b). Other genes that regulate CH size, such as *Lhx2*, are expressed in the cortical neuroepithelium rather than CH itself. *Lhx2* confers cortical identity in the telencephalic neuroepithelium, cell-autonomously suppressing CH fate (Mangale et al., 2008; Monuki et al., 2001).

LIM-homeodomain transcription factor Lmx1a is one of the earliest markers of the CH. In contrast to the aforementioned genes, its expression in the telencephalon is limited to dorsal midline structures – the CH and non-neural choroid plexus epithelium (ChPe) (Caronia-Brown et al., 2016; Chizhikov et al., 2019; Failli et al., 2002). Previously, we found that loss of *Lmx1a* results in an aberrant contribution of the *Lmx1a*-lineage cells to the hippocampus, suggesting a function of this gene in establishing the CH/hippocampus boundary (Chizhikov et al., 2010).

This phenotype, however, was traced to embryonic day (e) 10.5, when the dorsal midline has not yet fully differentiated into the ChPe and CH. Thus, it was unknown whether Lmx1a has a role in the CH development or its function is limited to the segregation of the dorsal midline lineage from the cortical neuroepithelium before the formation of the CH.

In this paper, we investigated the role of *Lmx1a* in CH development by performing *Lmx1a* loss- and gain-of-function studies, complemented by RNAseq, *in utero* electroporation, and genetic experiments to identify downstream mediators of *Lmx1a* function in CH. We found that loss of *Lmx1a* compromises expression of numerous CH and CR cell markers, expression of Wnt signaling molecules, and cell cycle exit/differentiation and migration of CR cells, resulting in specific hippocampal abnormalities. Overexpression of *Lmx1a* was sufficient to induce key CH features (*Wnt3a* expression, CR cells, downregulation of the cortical selector gene *Lhx2*) in the cortical neuroepithelium, introducing *Lmx1a* as a master regulator of CH development.

## Results

### *Lmx1a -/-* mice exhibit a compromised DG associated with proliferation, glial scaffold, and migration abnormalities

Although *Lmx1a* is not expressed in the hippocampus (Chizhikov et al., 2019; Chizhikov et al., 2010), development of the hippocampus depends on CH (Hevner, 2016; Subramanian and Tole, 2009a) and, thus, can serve as a “readout” of CH functioning in *Lmx1a-/-* mice. Extending previous histological studies (Kuwamura et al., 2005; Sekiguchi et al., 1992), we found a modest (~17%) reduction in the length of the hippocampal CA1-3 fields (Fig. 1A-C), but a dramatic reduction of the HF that overlays the DG (~55% reduction, Fig. 1A, B, D) and the number of DG (Prox1+) neurons (~78% reduction, Fig. 1D-G) in adult (postnatal day 21, P21) *Lmx1a-/-* mice. Input resistance, a major determinant of neuronal excitability (Yang et al., 2021), was aberrantly low in DG neurons in *Lmx1a-/-* mice (Fig. 1H-J). In P3 *Lmx1a-/-* mutants, Ctip2 immunohistochemistry (Roy et al., 2019) revealed intact patterning of the hippocampal subfields (Suppl. Fig. 1A, B), but already reduced number of DG neurons (Suppl. Fig. 1C-E). Thus, to identify the mechanisms mediating DG abnormalities in *Lmx1a-/-* mutants, we analyzed embryonic stages, focusing on processes known to shape the DG.

**Fig. 1.**
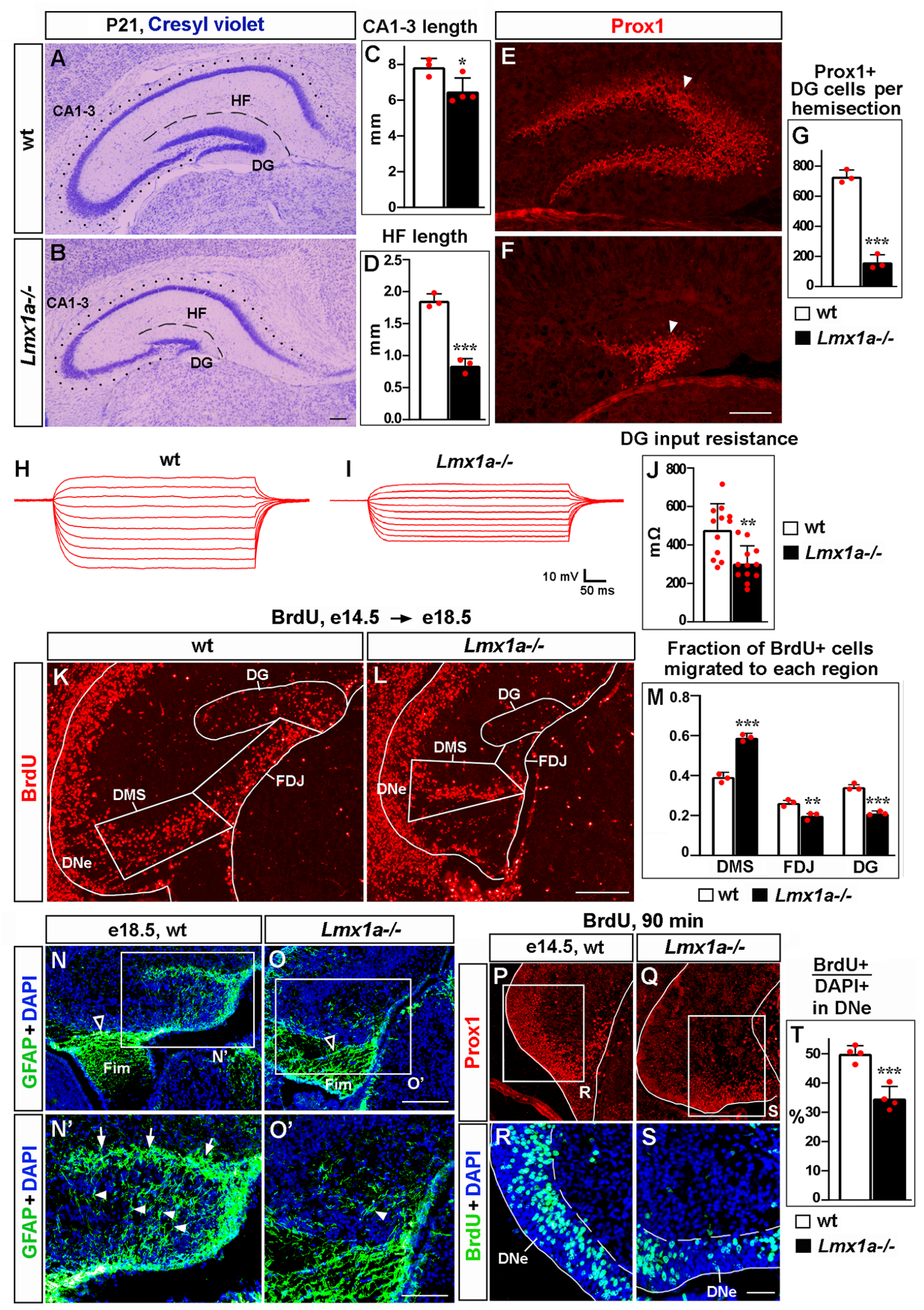
Loss of *Lmx1a* compromises DG development. In this and other figures, panels show coronal sections immunostained with indicated antibodies (unless noted otherwise). High magnification images show regions boxed in adjacent panels or diagrams. wt – wild type controls. (A-G) In P21 *Lmx1a-/-* mice, the length of the CA1-CA3 hippocampal fields (dotted line in A, B), length of the HF that overlays the DG (dashed line in A, B), and the number of Prox1+ DG neurons (arrowhead in E, F) were reduced (C, D, G). *p<0.05, ***p<0.001. n=3-4 animals per genotype. (H-J) Current-voltage curves (H, I) and a bar graph (J) showing a reduced input resistance of DG granule neurons in P21 *Lmx1a-/-* mice. **p<0.01, N=12 DG neurons from n=5 mice per genotype. (K-M) Mice were injected with BrdU at e14.5 to label progenitors in the DNe and analyzed at e18.5, when cells migrated toward the DG. A larger fraction of BrdU+ cells were located at the beginning of their migratory route (labeled as DMS, dorsal migratory stream, ***p<0.001) and smaller fractions of them reached the FDJ (**p<0.01) and DG (***p<0.001) areas in *Lmx1a-/-* mice. n=3 mice per genotype. (N-O’) By late embryogenesis, CH transforms into fimbria (Fim). The GFAP+ fimbrial scaffold (open arrowheads) was still present (N, O), but the transhilar scaffold was severely diminished in *Lmx1a-/-* mice (N’, O’). Arrowheads and arrows indicate GFAP+ glial fibers that cross the hilus and enrich at the HF, respectively. (P-T) Proliferation (% of BrdU+ cells among DAPI+ cells after a 90-min BrdU pulse) was reduced in the DNe (identified by Prox1 immunostaining) of *Lmx1a-/-* embryos. ***p<0.001, n=4 embryos per genotype. Scale bars: 250 μm (A, B, E, F, N, O); 200 μm (K, L); 100 μm (N’, O’, P, Q); 60 μm (R, S).

At e14.5, DG progenitors exit the dentate neuroepithelium (DNe) in the ventricular zone, forming the dentate migratory stream (DMS) and migrating to the fimbria-dentate junction (FDJ) and then to the DG (Nelson et al., 2020). To study cell migration, we labeled progenitors in the DNe by BrdU at e14.5 and analyzed embryos at e18.5 (Cai et al., 2018). Compared to controls, in *Lmx1a* mutants, a larger fraction of BrdU+ cells were located at the beginning of their migratory route (DMS), and fewer of them reached the FDJ and DG, revealing a compromised migration (Fig. 1K-M). Fimbrial and transhilar glial scaffolds are necessary for the migration of progenitors from the DNe to DG (Caramello et al., 2021; Hevner, 2016; Hodge et al., 2013). While the fimbrial scaffold appeared grossly normal, the transhilar scaffold was diminished in *Lmx1a* mutants (Fig. 1N-O’). A short (90-min) BrdU pulse, used to label cells in the S phase of the cell cycle (Lavado et al., 2013), revealed a reduced proliferation in the DNe of e14.5 *Lmx1a-/-* mutants (Fig. 1P-T). Thus, both proliferation and migration abnormalities contribute to DG deficits in *Lmx1a-/-* mice.

### Reduced exit of CH progenitors from the cell cycle and compromised migration of CH-derived CR cells in *Lmx1a-/-* mice

Next, we analyzed CH, where *Lmx1a* is expressed during embryogenesis. A short BrdU pulse and anti-activated Caspase 3 immunohistochemistry (Ivaniutsin et al., 2009; Miquelajauregui et al., 2007) revealed normal proliferation and apoptosis in the CH of *Lmx1a* mutants (Suppl. Fig. 2). In contrast, BrdU/Ki67 immunohistochemistry after a 24-hr BrdU pulse (Chizhikov et al., 2019) revealed a smaller fraction of progenitors exiting the cell cycle in *Lmx1a-/-* embryos (Fig. 2A-C), suggesting that *Lmx1a* is required for normal differentiation of CH progenitors.

**Fig. 2.**
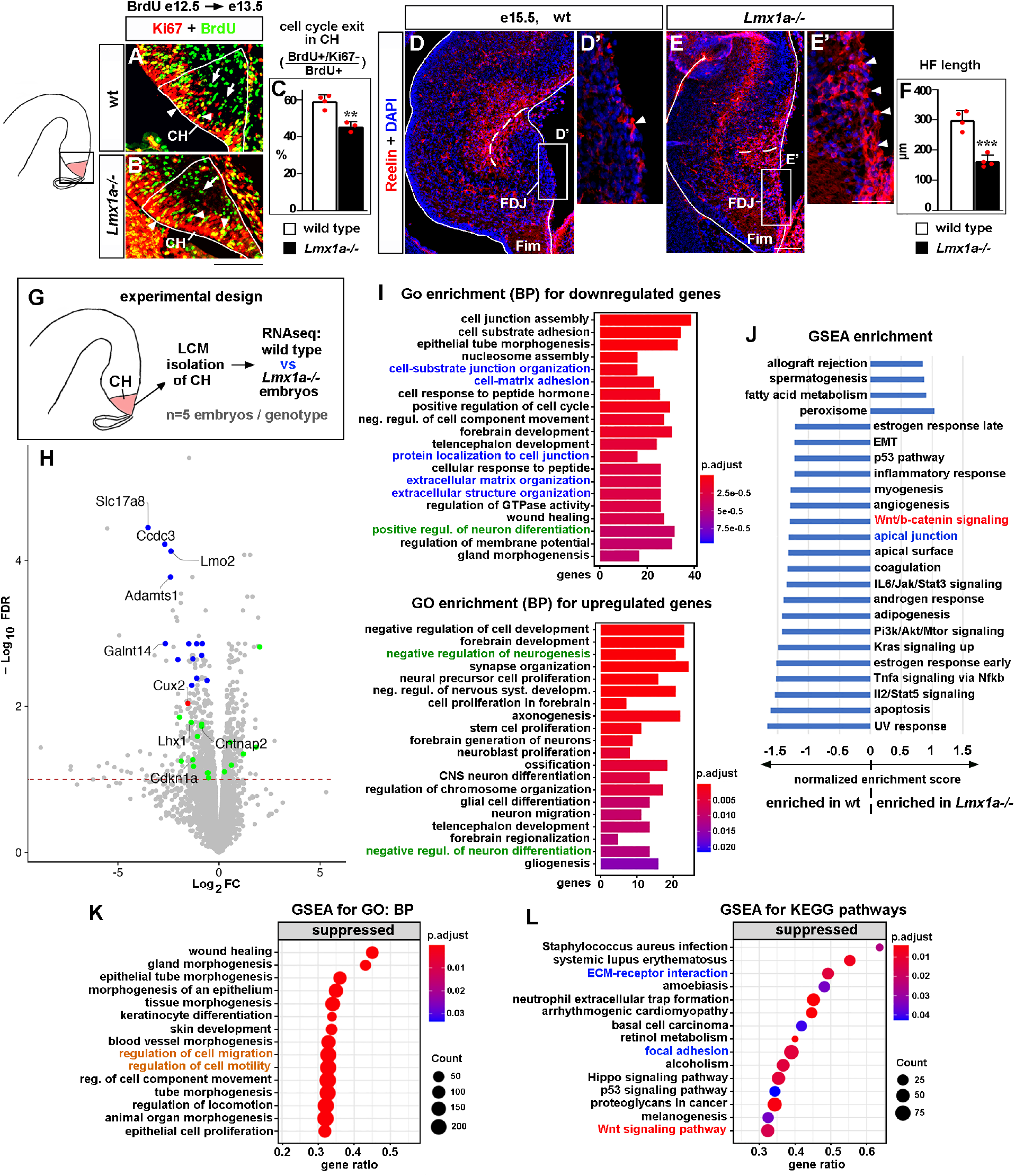
Developmental processes compromised in the CH of *Lmx1a-/-* mice. (A-C) A reduced fraction (%) of progenitors exited the cell cycle in the CH of *Lmx1a-/-* embryos. 24 h after BrdU injection, progenitors that exited the cell cycle were BrdU+/Ki67- (green, arrows); progenitors that re-entered the cell cycle were BrdU+/Ki67+ (yellow, arrowheads). **p<0.01, n=3-4 embryos per genotype. (D-F) By e15.5, in wild type embryos, many CR cells have already migrated into the HF area, and few CR cells remain at the FDJ surface (D, D’, arrowhead). In contrast, many CR cells were still located at the FDJ surface in *Lmx1a-/-* littermates (E, E’, arrowheads), which was associated with a reduced HF length (dashed line in D, E; F). ***p<0.001, n=4 embryos per genotype. (G) Experimental design for RNAseq analysis. (H) Volcano plot of transcripts detected by RNAseq. Transcripts above the dashed line (FDR=0.1) were considered differentially expressed in *Lmx1a-/-* CH. CH markers, identified among the 100 most misregulated genes, are shown in blue (also see Suppl. Fig. 4); genes associated with CR cells (Franzén et al., 2019; Li et al., 2021) are shown in green; the only previously identified *Lmx1a* target in the CH, *Cux2* (Fregoso et al., 2019), is shown in red. (I) GO (biological process, BP) enrichment analysis for the genes differentially expressed in *Lmx1a-/-* CH. (J) Pathways enriched in wild type and *Lmx1a-/-* CH based on GSEA analysis of RNAseq data. (K, L) Top 15 pathways/processes suppressed in the *Lmx1a-/-* CH based on GSEA for GO (BP) and GSEA for KEGG analyses. Processes/pathways related to neuronal differentiation, cell migration, Wnt signaling, and tissue integrity are highlighted in green, orange, red, and blue, respectively. Scale bars: 100 μm (A, B, D, E); 50 μm (D’, E’).

Since CH progenitors differentiate into CR cells (Dixit et al., 2011; Gu et al., 2011), we studied CR cells. After exiting the CH, CR cells first migrate tangentially along the FDJ and then **–** radially into the hippocampal field to form the hippocampal fissure (HF) (Causeret et al., 2021). Immunohistochemistry against the CR cell marker Reelin (Hodge et al., 2013; Ledonne et al., 2016; Siegenthaler and Miller, 2008) revealed that in e16.5 wild type embryos, many CR cells have populated the HF, and few remained at the FDJ. In contrast, in *Lmx1a-/-* littermates, many CR cells were still at the FDJ and the HF was abnormally short (Fig. 2D-F). CR cell distribution and HF abnormalities persisted at P3 (Suppl. Fig. 3), indicating that loss of *Lmx1a* leads to severe long-lasting deficits in the migration of CR cells.

### Genes and developmental processes misregulated in the CH of *Lmx1a-/-* mice based on RNAseq analysis

To identify CH genes regulated by *Lmx1a*, we performed RNAseq analysis. For these experiments, we isolated CH by laser capture microdissection (LCM) from wild type and *Lmx1a -/-* embryos at e13.5, when the hippocampus begins to develop, and the CR cells that populate the HF emerge from the CH (Gu et al., 2011; Nelson et al., 2020). We found that 612 genes were significantly downregulated, and 457 genes were significantly upregulated in the CH of *Lmx1a* mutants compared to wild type littermates (Fig. 2G, H). Notably, these genes included *Cux2* (Fig. 2H, red dot), the only previously known *Lmx1a* downstream gene in the CH (Fregoso et al., 2019), validating our experimental approach.

Of the 100 genes most misregulated in *Lmx1a-/-* CH (FDR<0.05), 55 had *in situ* hybridization images in public databases (Gene Paint, Eurexpress, Allen Brain Atlas). Interestingly, 14 of these genes (~25%) were specifically or predominantly expressed in the CH and were downregulated in *Lmx1a* mutants (Fig. 2H, blue dots, and Suppl. Fig. 4), including a previously described CH marker *Lmo2* (Mangale et al., 2008) and 13 novel CH markers (*Slc17a8, Ccdc3, Adamts1, Galnt14, Camk2a, Arfgef3, Rdh10, Plxnc1, Kdr, Asb4, Slc14a2, Peg3, Slit3*). Functionally, these genes varied considerably, ranging from transcription factors (*Lmo2*) to transporters of molecules (*Slc17a8, Slc14a2*) to lipid and carbohydrate metabolism regulators (*Ccdc3, Rhd10*) to transmembrane receptors (*Plxnc1*).

Having observed CR cell abnormalities in *Lmx1a* mutants (Fig. 2D-E’), we studied whether our newly identified Lmx1a downstream targets include genes associated with CR cells, taking advantage of recently identified datasets of genes enriched in CR cells (Franzén et al., 2019; Li et al., 2021). 11 known CR cell-enriched genes were downregulated (*Car10, Lhx1, Trp73, Cntnap2, Reln, Cdkn1a, Chst8, Spock1, Akap6, Cacna2d2, Tbr2*) and six – were upregulated (*Tmem163, Sema6a, Lhx5, Lingo1, Edil3, MAPK8*) in the CH of *Lmx1a-/-* embryos (green dots in Fig. 2H). Thus, loss of *Lmx1a* diminishes the expression of a diverse set of CH markers and compromises the expression of CR cell-associated genes in the CH.

To further test for an intrinsic role of *Lmx1a* in the CH, we *in utero* electroporated the CH of e11 wild type embryos with either previously validated anti-*Lmx1a* shRNA (Fregoso et al., 2019) or control shRNA together with GFP. At e13.5, GFP+ cells in the CH were isolated by LCM, and gene expression was analyzed by qRT-PCR (Suppl. Fig. 5). In these experiments, we confirmed the downregulation of all seven tested Lmx1a downstream targets, identified by RNAseq, including both CH and CR markers/key developmental genes (see below) (Suppl. Fig. 5), supporting an intrinsic function of *Lmx1a* in the CH.

To gain further insight into the developmental processes and pathways affected in the CH by *Lmx1a* loss, we performed bioinformatics analysis of our RNAseq data (Fig. 2I-L). Interestingly and consistently with a reduced cell cycle exit of CH progenitors (Fig. 2A-C), Gene Ontology (GO) enrichment analysis revealed that the “positive regulation of neuronal differentiation” category was overrepresented in genes significantly downregulated, while the “negative regulation of neurogenesis” and “negative regulation of neuronal differentiation” categories were overrepresented in genes significantly upregulated in the CH of *Lmx1a-/-* embryos (Fig. 2I, highlighted in green). Consistent with the abnormal distribution of CR cells (Fig. 2D-E’), Gene Set Enrichment Analysis (GSEA) for GO revealed that “regulation of cell migration” and “regulation of cell motility” processes were suppressed in *Lmx1a-/-* mutants (Fig. 2K, highlighted in orange). In addition, the GSEA revealed enrichment of Wnt/β-catenin signaling in wild type relative to *Lmx1a-/-* CH (Fig. 2J). Similarly, GSEA for Kyoto Encyclopedia of Genes and Genomes (KEGG) revealed suppressed Wnt signaling in the CH of *Lmx1a-/-* mice (Fig 2L, highlighted in red). Finally, GO enrichment, GSEA, and GSEA for KEGG analyses predicted compromised cell-cell and cell-extracellular matrix adhesion in *Lmx1a-/-* CH (Fig. 2I, J, L, related processes are highlighted in blue), suggesting that abnormal cell adhesion contributes to previously reported aberrant dispersion of *Lmx1a*-lineage cells into the adjacent hippocampus in *Lmx1a* mutants (Chizhikov et al., 2010). Thus, *Lmx1a* is required for normal expression of diverse CH markers and CR cell genes, and our bioinformatics analysis supports the role of *Lmx1a* in the differentiation and migration of CR cells, and canonical Wnt signaling.

### *Lmx1a* regulates the number and input resistance of DG neurons, proliferation of DG progenitors, and the transhilar glial scaffold via secreted Wnt3a

Canonical Wnt signaling promotes proliferation in the DNe and transhilar scaffold formation (Li and Pleasure, 2005; Zhou et al., 2004), suggesting that *Lmx1a* regulates DG morphogenesis at least partially via secreted Wnts produced in the CH. Our RNAseq analysis revealed that two Wnts known to activate canonical Wnt signaling, Wnt2b and Wnt9a, both expressed specifically in the CH, were significantly (FDR<0.05) downregulated in *Lmx1a-/-* mutants (Suppl. Fig. 6). *Wnt2b* knockout mice have a normally-sized DG (Tsukiyama and Yamaguchi, 2012), while Wnt9a, although capable of activating canonical Wnt signaling in the liver (Matsumoto et al., 2008), primarily acts as non-canonical Wnt ligand (Nie et al., 2020). In contrast, the knockout of CH-specific *Wnt3a* resulted in a nearly absent DG (Lee et al., 2000). Our RNAseq analysis showed that *Wnt3a* was downregulated in the CH of e13.5 *Lmx1a-/-* mice, although the difference did not reach a statistical significance (FDR=0.13, Suppl. Fig. 6). However, quantification of *in situ* hybridization signal, revealed a significant (p<0.01) downregulation of *Wnt3a* expression in the CH of *Lmx1a-/-* embryos at e14 (Fig. 3A-C), just before we detected a reduced proliferation in the DNe (Fig. 1P-T). Knockdown of *Lmx1a* in the CH resulted in a reduced expression of *Wnt3a* (Suppl. Fig. 5I), supporting an intrinsic role for *Lmx1a* in regulating *Wnt3a* expression.

**Fig. 3.**
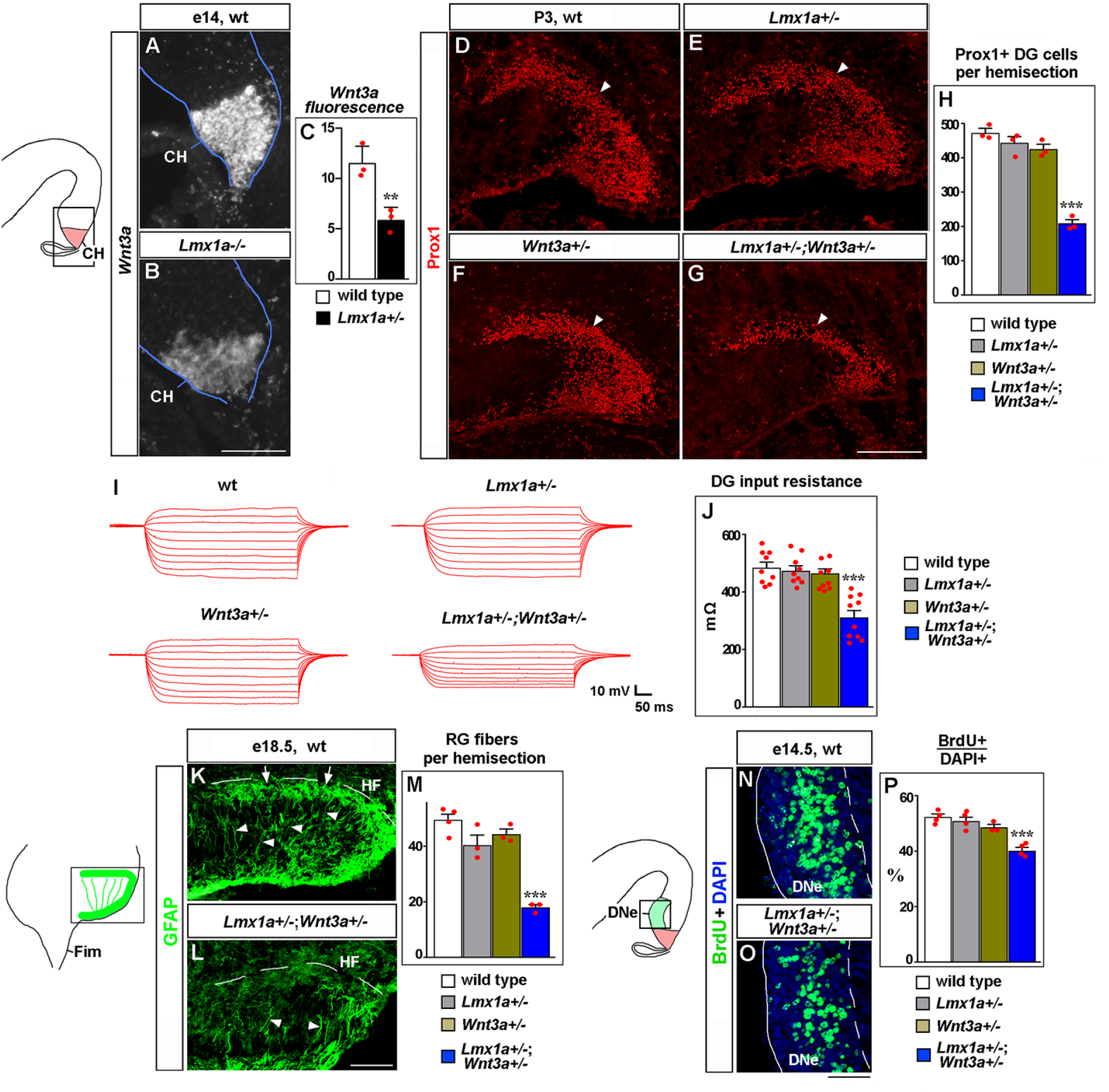
*Lmx1a* regulates DG development via *Wnt3a*. (A-C) The intensity of *Wnt3a in situ* hybridization signal (white) in the CH was reduced in e14 *Lmx1a-/-* embryos. **p<0.01, n=3 embryos per genotype. (D-H) The number of Prox1+ DG neurons (arrowhead) was reduced in *Lmx1a+/-;Wnt3a+/-* double heterozygotes, but not single-gene heterozygotes, compared to wild type controls at P3. ***p<0.001, n=3 mice per genotype. (I, J) Current-voltage curves (I) and a bar graph (J) showing a reduced input resistance of DG granule neurons in *Lmx1a+/-;Wnt3a+/-* double heterozygotes, but not single-gene heterozygotes, compared to wild type controls at P21. ***p<0.001, N=9-10 DG neurons from n=4-5 mice per genotype. (K-M) The number of GFAP+ transhilar glial fibers was reduced in *Lmx1a+/-;Wnt3a+/-* double heterozygotes, but not single-gene heterozygotes, compared to wild type controls. ***p<0.001, n=3-4 mice per genotype. Arrowheads and arrows indicate fibers that cross the hilus and enrich at the HF, respectively. (N-P) Proliferation (% of BrdU+ cells after a 90-min BrdU pulse) in DNe was reduced in *Lmx1a+/-;Wnt3a+/-* double heterozygotes, but not single-gene heterozygotes, compared to wild type controls. ***p<0.001, n=3-4 mice per genotype. Scale bars: 100 μm (A, B, K, L); 200 μm (D-G); 50 μm (N, O).

Since normal DG development is particularly sensitive to the level and dynamics of canonical Wnt signaling (Arredondo et al., 2020; Zhou et al., 2004), to test Wnt3a as a downstream mediator of Lmx1a function in CH/DG development, we performed an analysis of *Lmx1a/Wnt3a* double heterozygotes rather than *Wnt3a* overexpression rescue experiments in *Lmx1a -/-* mice. In contrast to wild type or single-gene heterozygous (*Lmx1a+/-* or *Wnt3a+/-*) mice, their double heterozygous *Lmx1a+/-;Wnt3a+/-* littermates exhibited a reduced number (Fig. 3D-H) and input resistance (Fig. 3I, J) of DG neurons, and transhilar glial scaffold abnormalities (Fig. 3K-M). A reduced number of DG neurons was associated with reduced proliferation in DNe (Fig. 3N-P). Together, these data provide evidence that Lmx1a and Wnt3a act in the same pathway to regulate DG morphogenesis.

### *Lmx1a* promotes migration of CR cells and the HF and transhilar scaffold formation via *Tbr2*

CR cells promote HF and radial glial scaffold formation (Frotscher et al., 2003; Hodge et al., 2013; Meyer et al., 2004; Meyer et al., 2019). Having observed CR cell migration deficits in *Lmx1a -/-* mutants (Fig. 2D-E’), we studied *Lmx1a*-dependent mechanisms of CR cell migration. For this purpose, we searched the literature to find whether mutants for CR cell-associated genes misregulated in *Lmx1a-/-* CH (Fig. 2H, green dots) have phenotypes similar to those in *Lmx1a-/-* mutants. Interestingly, *Nestin-Cre;Tbr2^floxed/floxed (F/F)^* mice exhibited CR cell migration, HF, and glial scaffold abnormalities (Hodge et al., 2013) that resemble those in *Lmx1a-/-* mice. While in the CH, *Tbr2* is expressed specifically in differentiating CR cells, this gene is also expressed in intermediate hippocampal (DG) progenitors. *Nestin-Cre* mice delete floxed sequences in both the CH and hippocampal progenitors (Hodge et al., 2013), making it difficult to identify the developmental origin of the CR/ hippocampal phenotypes in *Nestin-Cre:Tbr2^F/F^* mice.

To study the intrinsic role of *Tbr2* in CR cells, we generated and analyzed *Lmx1a-Cre;Tbr2^F/F^* mice, in which *Tbr2* was specifically inactivated in the CH but still expressed in hippocampal progenitors (Suppl. Fig. 7A-C’). Similar to *Nestin-Cre;Tbr2^F/F^* mice (Hodge et al., 2013), *Lmx1a-Cre;Tbr2^F/F^* mice exhibited aberrant accumulation of Reelin+ CR cells at the FDJ, associated with a reduced HF length (Suppl. Fig. 7D-F), indicating that *Tbr2* acts intrinsically in CR cells to regulate their migration and HF/DG morphogenesis.

Consistent with our RNAseq data (Fig. 4A), immunohistochemistry revealed reduced Tbr2 expression specifically in the CH of *Lmx1a-/-* mice at e13 (Fig. 4B-D), when CR cells that predominantly populate the HF emerge from the CH (Gu et al., 2011). Knockdown of *Lmx1a* in the CH also resulted in reduced expression of *Tbr2* (Suppl. Fig. 5J), supporting an intrinsic role for *Lmx1a* in the regulation of *Tbr2* expression. Since CR cell migration and DG morphogenesis are complex processes that require precise expression levels of key genes (Gil et al., 2014; Ha et al., 2020; Hevner, 2016), to study whether *Tbr2* is a downstream mediator of *Lmx1a* function in CR cells/DG development, we performed an analysis of *Lmx1a/Tbr2* double heterozygotes rather than *Tbr2* overexpression rescue experiments in *Lmx1a -/-* mice. In contrast to wild type controls or single-gene heterozygotes (*Lmx1a+/-* or *Lmx1a-Cre;Tbr2^+/F^*, in which one copy of *Tbr2* was specifically deleted in CH), their *Lmx1a+/-;Lmx1a-Cre;Tbr2^+/F^* double-gene heterozygous littermates revealed aberrant accumulation of Reelin+ CR cells at the FDJ, reduced HF length and transhilar scaffold abnormalities (Fig. 4E-F, Suppl. Fig. 8). These phenotypes were not associated with a delayed exit of CH progenitors from the cell cycle (Fig. 4G-H). Thus, *Lmx1a* acts, at least partially, via *Tbr2* to regulate migration of CR cells and HF/transhilar glial scaffold formation.

**Fig. 4.**
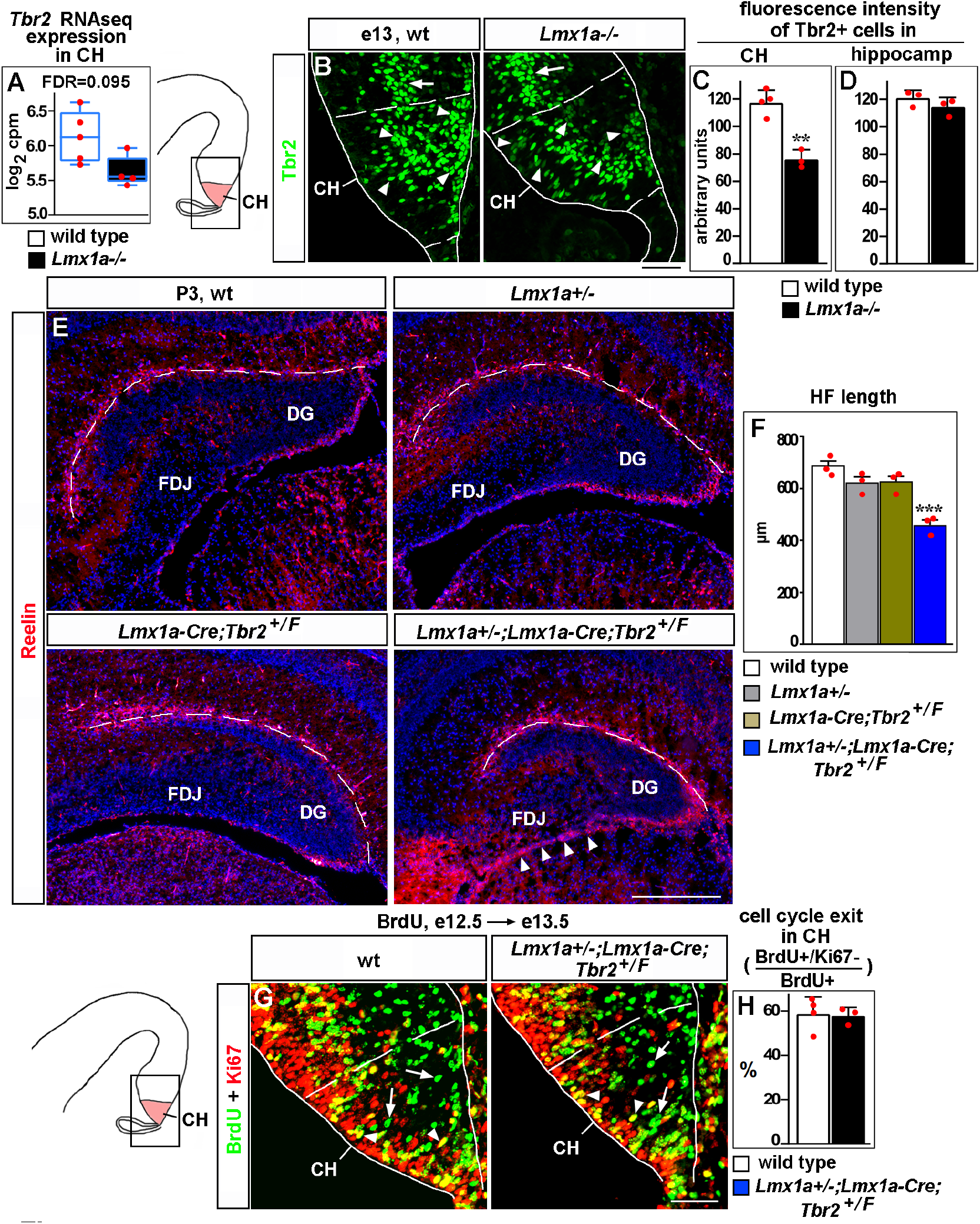
*Lmx1a* regulates CR cell migration and HF formation via *Tbr2*. (A) Normalized read counts for *Tbr2* from the RNAseq experiment (Fig. 2G, H). (B-D) Arrowheads indicate Tbr2+ cells in the CH (CR cells, Hodge et al., 2013) (B), which exhibited lower intensity of Tbr2 immunofluorescence in e13 *Lmx1a-/-* embryos compared to controls (B, C). In contrast, Tbr2+ cells in the hippocampal primordium (hippocampal intermediate progenitors, Hodge et al., 2013) (arrow in B) had similar Tbr2 immunofluorescence intensity in control and *Lmx1a-/-* embryos (B, D), suggesting that *Lmx1a* loss reduces *Tbr2* expression specifically in cells arising in the CH. (Dashed lines demarcate the CH boundaries, identified by Lmx1a immunostaining of adjacent sections, as described in the Materials and Methods). (E-F) Reduced HF length (dashed line) and aberrant accumulation of CR cells at the FDJ surface (arrowheads) in *Lmx1a/Tbr2* double-gene heterozygotes (*Lmx1a+/-;Lmx1a-Cre;Tbr2^+ F^* mice – *Lmx1a+/-* mice in which one copy of *Tbr2* was deleted specifically in the CH), but not singlegene heterozygotes, compared to wild type controls at P3. ***p<0.001, n=3 mice per genotype. (G, H) Normal cell cycle exit of progenitors in the CH of *Lmx1a/Tbr2* double heterozygous embryos. 24 h after BrdU injection, progenitors that exited the cell cycle were BrdU+/Ki67-(green, arrows); progenitors that re-entered the cell cycle were BrdU+/Ki67+ (yellow, arrowheads). n=3-4 mice per genotype. Scale bars: 50 μm (B, G); 200 μm (E).

### *Lmx1a* promotes cell cycle exit and differentiation of CR cells in the CH via *Cdkn1a*

One of the CR markers misregulated in the *Lmx1a-/-* CH (Fig. 2H) was *Cdkn1a*, the gene known to promote exit from the cell cycle in several non-neural and neural cell types (Xiao et al., 2020). Consistent with our RNAseq analysis (Fig. 5A), immunohistochemistry revealed reduced *Cdkn1a* expression in the CH of *Lmx1a-/-* mice at e13 (Fig. 5B-D). Knockdown of *Lmx1a* in the CH resulted in a reduced expression of *Cdkn1a* (Suppl. Fig. 5K), supporting an intrinsic role for *Lmx1a* in regulating *Cdkn1a* expression. To study whether decreased *Cdkn1a* expression mediates a reduced cell cycle exit of CH progenitors in *Lmx1a-/-* embryos (Fig. 2A-C), we performed rescue experiments, *in utero* electroporating *Cdkn1a+GFP* or *GFP* alone (control) into the CH. In these experiments, plasmids are taken up by Ki67+ progenitor cells that line the lateral ventricles (Lavado et al., 2013), into which the plasmids are injected. Expression of exogenous *Cdkn1a* in the CH of *Lmx1a-/-* embryos increased the fraction of progenitors that exited the cell cycle (GFP+/Ki67-cells among GFP+ cells), making it comparable to that in controls (CH of wild type embryos electroporated with *GFP* alone) (Fig. 5E-H). Interestingly, *Cdkn1a* electroporation also normalized the number of differentiating (p73+) (Meyer et al., 2004; Siegenthaler and Miller, 2008) CR cells in *Lmx1a-/-* embryos (Fig. 5I-L), indicating that decreased *Cdkn1a* expression contributes to both reduced cell cycle exit in CH and decreased differentiation of CR cells in *Lmx1a-/-* embryos.

**Fig. 5.**
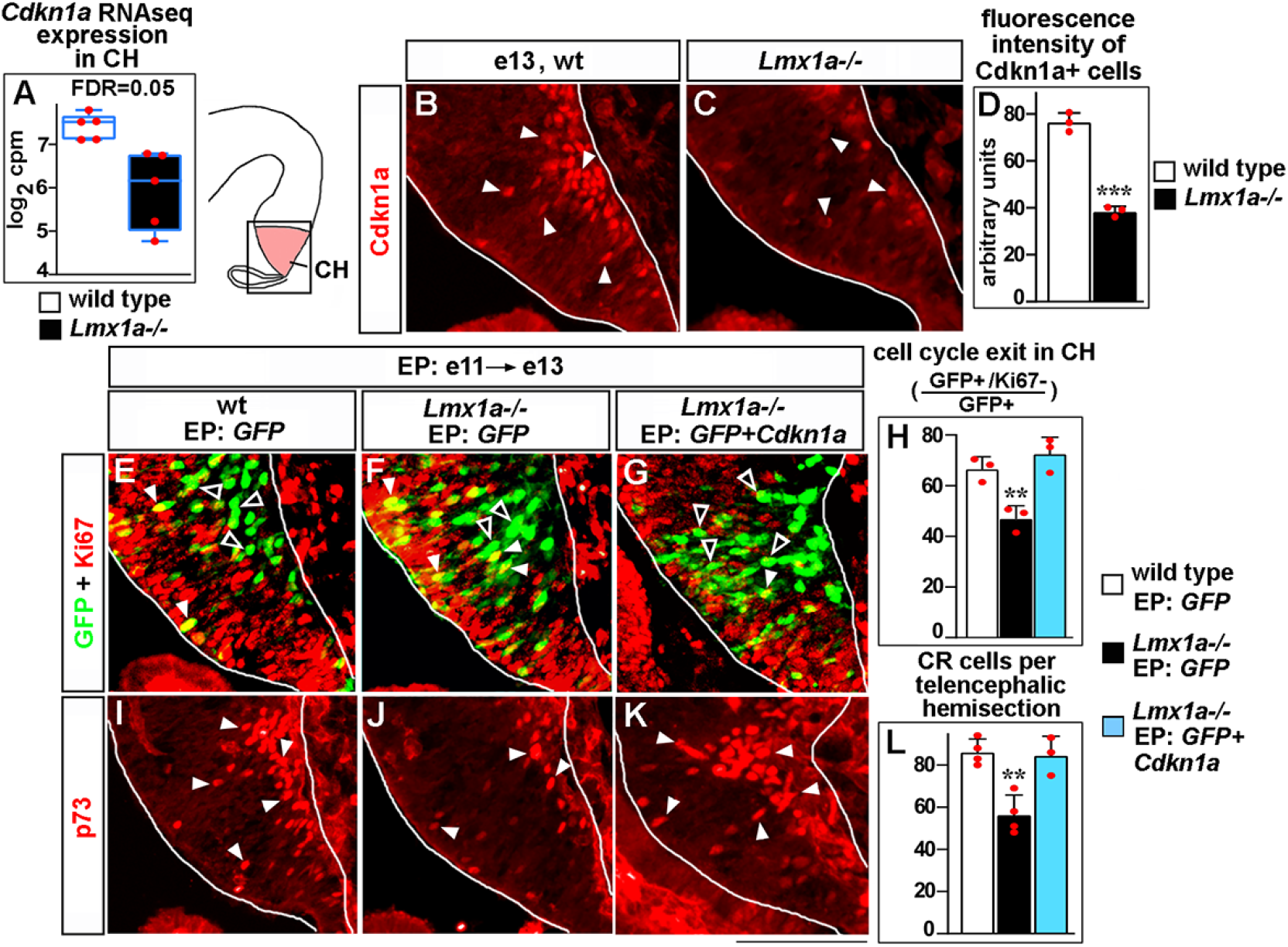
*Lmx1a* regulates exit of progenitors from the cell cycle and CR cell differentiation in the CH via *Cdkn1a*. (A) Normalized read counts for *Cdkn1a* from the RNAseq experiment (Fig. 2G, H). (B-D) Arrowheads indicate Cdkn1a+ cells in the CH (B, C), which exhibited lower immunofluorescence intensity in e13 *Lmx1a-/-* embryos (B-D). ***p<0.001, n=3 embryos per genotype. (E-L) Embryos were *in utero* electroporated (EP) with indicated genes at e11 and analyzed at e13. Sections in panels E, F, and G are adjacent to those in panels I, J, and K, respectively. (E-G) Arrowheads indicate electroporated cells that re-entered the cell cycle (GFP+/Ki67+ cells, yellow); open arrowheads indicate electroporated cells that exited the cell cycle (GFP+/Ki67-cells, green). The reduced exit of progenitors from the cell cycle in *Lmx1a-/-* CH was rescued by electroporation of *Cdkn1a* (H). **p<0.01 versus *GFP* electroporated wild type and *Cdkn1a+GFP* electroporated *Lmx1a-/-* embryos. n=3 embryos per condition. (I-L) Arrowheads indicate p73+ CR cells. The reduced number of CR cells in *Lmx1a-/-* mutants was rescued by electroporation of *Cdkn1a* (L). **p<0.01 versus *GFP* electroporated wild type and *Cdkn1a+GFP* electroporated *Lmx1a-/-* embryos. n=3-4 embryos per condition. Scale bar: 100 μm.

### *Lmx1a* is sufficient to induce ectopic CH

Having identified the role of *Lmx1a* in multiple aspects of CH development, we tested whether its expression alone is sufficient to confer CH fate in the telencephalon. We *in utero* electroporated *Lmx1a* into the medial telencephalon of e10.75 wild type embryos, after the cortical neuroepithelium had already been specified by *Lhx2* (Mangale et al., 2008), and analyzed three key cortical hem parameters: 1) expression of the CH signaling molecule Wnt3a (Grove et al., 1998), 2) the appearance of ectopic p73+ CR cells (Meyer et al., 2004; Siegenthaler and Miller, 2008), and 3) the suppression of *Lhx2* expression, a key feature of native CH (Mangale et al., 2008).

Excitingly, in contrast to *GFP* controls, *Lmx1a+GFP* electroporation induced ectopic *Wnt3a*. Extensive overlap between the GFP and *Wnt3a* signals suggested cell-autonomous induction of *Wnt3a* by *Lmx1a* in these experiments (Fig. 6A-D’). Normally, CR cells, identified by the expression of their specific marker p73, arise deep in the CH and then migrate along the outer surface of the telencephalic neuroepithelium (Meyer et al., 2004; Siegenthaler and Miller, 2008). In contrast to *GFP* controls, in *Lmx1a*-electroporated samples, we observed numerous ectopic p73+ cells deep in the neuroepithelium beyond the CH (Fig. 6E-H’), which was associated with a dramatic increase in the total number of p73+ cells (Fig 6E-I). Finally, quantification of the immunohistochemistry signal revealed a significant reduction of Lhx2 expression in hippocampal primordium cells electroporated with *Lmx1a+GFP* versus those electroporated with *GFP* alone (Fig. 6J-L). Thus, *Lmx1a* expression is sufficient to induce key CH features in the medial telencephalic neuroepithelium.

**Fig. 6.**
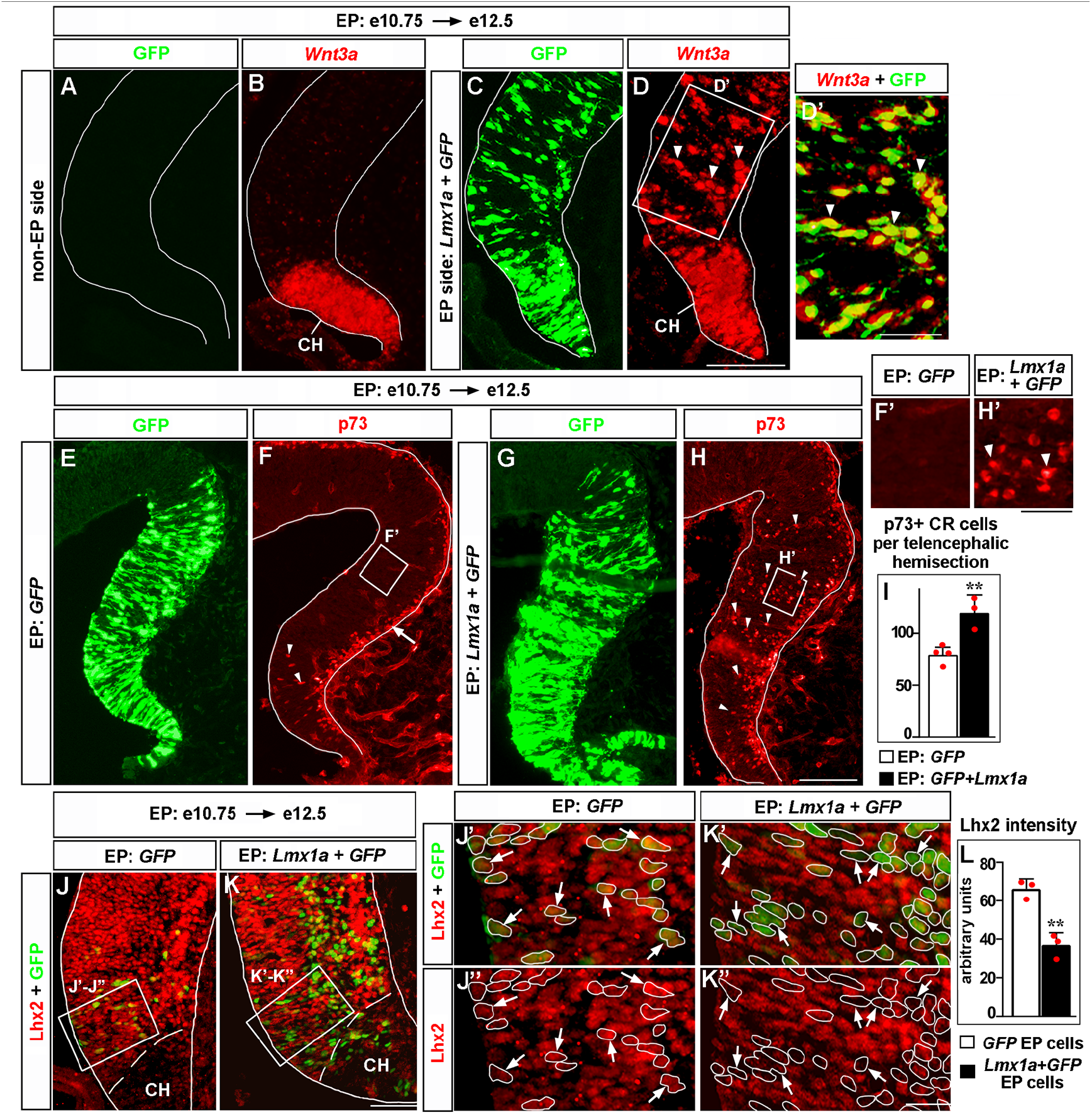
*Lmx1a* is sufficient to induce key features of the CH. Panels A and B, C and D, J’ and J’’, K’ and K’’ show the same sections imaged for different markers. Panels E and F, G and H show adjacent sections. wild type embryos were *in utero* electroporated (EP) at e10.75 and analyzed at e12.5. (A-D’) In contrast to the non-electroporated (non-EP) side, in which *Wnt3a* expression is limited to the CH (A, B), *Lmx1a* electroporation (EP side) induces ectopic *Wnt3a* expression in the hippocampal field (arrowheads), clearly beyond the CH (C, D). Extensive overlap between GFP fluorescence and *Wnt3a in situ* hybridization signal (D’, arrowheads) indicates cell-autonomous induction of *Wnt3a* by *Lmx1a*. (E-I) In *GFP*-electroporated controls (F, F’), p73+ CR cells were deeply located (arrowheads) only in the CH (where p73+ CR cells arise, Siegenthaler and Miller, 2008), while migrating superficially-located p73+ CR cells (arrow) were found also in the cortical neuroepithelium. In *Lmx1a*-electroporated embryos, p73+ cells were found deeply located (arrowheads) throughout the medial telencephalic neuroepithelium (H, H’). The total number of p73+ CR cells increased in *Lmx1a*-electroporated embryos (I) (**p<0.01, n=3-4 embryos per condition), further supporting induction of CR cells by *Lmx1a*. (J-L) *Lhx2* is expressed in the hippocampal primordium but not in the CH (J, K). Arrows indicate cells in the hippocampal primordium electroporated with *GFP* (controls) (J’, J’’). Arrowheads indicate such cells electroporated with *Lmx1a+GFP* (K’, K’’). The intensity of Lhx2 immunofluorescence was reduced in *Lmx1a*-electroporated cells (J’-L), indicating that *Lmx1a* is sufficient to repress *Lhx2* expression. **p<0.01, n=3 embryos per condition. Scale bars: 100μm (A-D, E-H); 50 μm (D’, J, K); 30 μm (F’, H’); 15 μm (J”-K’’).

## Discussion

While the critical role of signaling centers in the development of the vertebrate CNS has been recognized for decades, very little is known about how distinct cellular and molecular mechanisms are coordinated to achieve the proper formation and function of specific signaling centers (Bielen et al., 2017; Cavodeassi and Houart, 2012; Manfrin et al., 2019; Subramanian and Tole, 2009b). Here we show that *Lmx1a* acts as a master regulator of the CH, a major signaling center that controls the formation of the hippocampus in the medial telencephalon (Caronia-Brown et al., 2014; Mangale et al., 2008; Moore and Iulianella, 2021). Our loss-of-function experiments revealed that *Lmx1a* is required for the expression of a broad range of CH markers and Wnt signaling molecules, the proper exit of CH progenitors from the cell cycle, and differentiation and migration of CR cells. In complementary gain-of-function experiments, we found that *Lmx1a* is sufficient to induce key CH features (*Wnt3a* expression, CR cells, and downregulation of expression of the cortical selector gene *Lhx2*, involved in the CH/hippocampus boundary formation) (Fig. 7). This study identifies *Lmx1a* as the first intrinsic molecule that specifies CH fate and, to our knowledge, provides the strongest example that the orchestration of signaling center development in the CNS utilizes master regulatory genes.

**Fig. 7.**
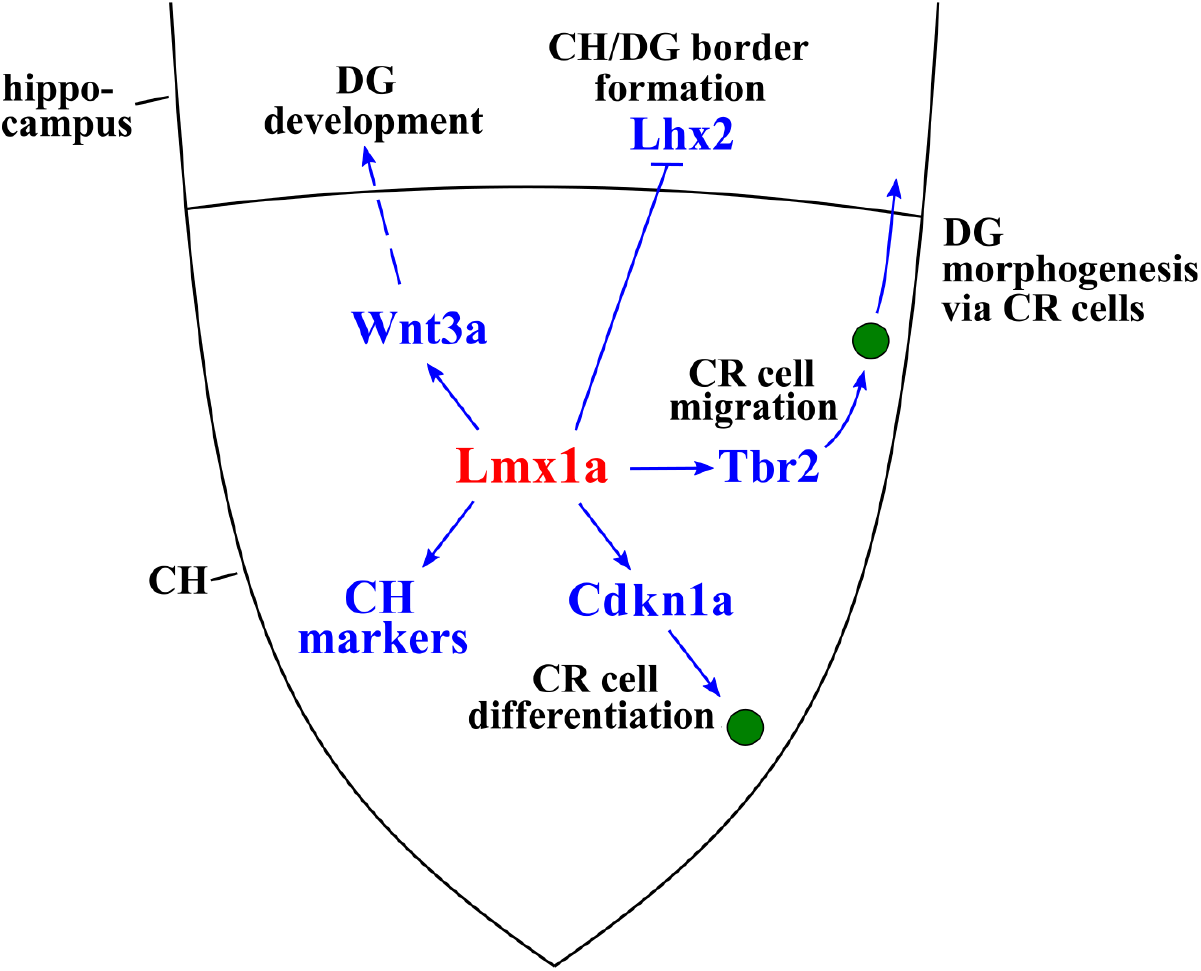
*Lmx1a*-dependent developmental processes and downstream mediators of *Lmx1a* activity in the CH. *Lmx1a* promotes expression (solid arrow) of secreted Wnt3a, which non-autonomously (dashed arrow) regulates the development of the hippocampal DG (proliferation in the DNe, the transhilar scaffold, and the number and input resistance of DG neurons). *Lmx1a* represses the expression of the cortical selector gene *Lhx2* (solid bar), segregating the CH from the adjacent hippocampal field. *Lmx1a* positively regulates the expression of a wide range of CH markers. It also promotes the exit of CH progenitors from the cell cycle and their differentiation into CR cells by activating the expression of *Cdkn1a*, and promotes migration of CR cells via *Tbr2*, which is necessary for the HF and transhilar scaffold formation.

Until now, no intrinsic molecule had been reported to induce CH. Although the loss of transcription factors *Gli3* and *Dmrt3/4/5* prevents CH formation, expression domains of all these genes expand beyond the CH, suggesting that they play permissive rather than instructive roles in CH development (Grove et al., 1998; Kikkawa and Osumi, 2021; Quinn et al., 2009; Subramanian et al., 2009a; Subramanian and Tole, 2009b). Furthermore, mosaic inactivation of the cortical selector gene *Lhx2*, prior to e10.5, resulted in the development of ectopic CH in *Lhx2*-null patches of the cortical neuroepithelium, suggesting that CH is the default fate in the medial telencephalon (Mangale et al., 2008). However, our overexpression of *Lmx1a* in the hippocampal primordium at e10.75, after the cortical neuroepithelium identity has already been assigned by *Lhx2* (Mangale et al., 2008), resulted in cell-autonomous ectopic expression of the key CH signaling molecule Wnt3a and the generation of ectopic p73+ CR cells, indicating that the CH identity is actively assigned by *Lmx1a*. Recently, *Lmx1a* was reported to activate a human CH-specific enhancer and an enhancer for the mouse *Cux2*, gene expressed in the CH and a subset of cortical neurons (Fregoso et al., 2019). Although the significance of the aforementioned enhancers for CH development or function remains unknown, these data are consistent with and further support the role of *Lmx1a* in the induction of the CH that we describe in the current study.

Interestingly, although knockout/knockdown of *Lmx1a* dramatically reduces the expression of numerous CH markers (Suppl. Figs. 5 and 6), the CH was not completely lost in *Lmx1a-/-* mice (Fig. 3A-C) (Chizhikov et al., 2010), suggesting a partial compensation for the *Lmx1a* CH-inducing activity by another gene. Ablation of the Bmp signaling (in *BmpR1a/b* double knockout mice) was reported to prevent CH induction (Fernandes et al., 2007), while mice with downregulated Bmp signaling still develop a CH-like structure at the telencephalic dorsal midline (Chizhikov et al., 2019; Hébert et al., 2002). Moreover, while high levels of Bmps promoted the ChPe fate (Doan et al., 2012; Watanabe et al., 2012), low Bmp doses induced CH gene expression signature in embryonic stem cells *in vitro* (Watanabe et al., 2016). Thus, it is likely that an intrinsic molecule(s) activated by low levels of Bmp signaling acts partially redundant to *Lmx1a* to assign CH fate in the medial telencephalic neuroepithelium.

In addition to CH induction, we report that *Lmx1a* functions as a regulator of expression of CH-specific Wnt signaling molecules, including the canonical Wnt3a (Fig. 3A-C, Suppl. Figs. 5I, 6). Canonical Wnt signaling from the CH promotes the proliferation of hippocampal (DNe) progenitors and the formation of the hippocampal radial glial scaffold that guides the migration of progenitors from the DNe to the DG (Li and Pleasure, 2005; Zhou et al., 2004). Our analysis of *Lmx1a+/-; Wnt3a+/-* double heterozygous mice confirmed that *Lmx1a* and *Wnt3a* act in the same pathway to regulate the proliferation of DG progenitors, radial glial scaffold formation, and electrophysiological properties (input resistance) of DG neurons.

We found that besides CH signaling, *Lmx1a* regulates two distinct steps of CR cell development – their migration and differentiation from CH progenitors. Migrating CR cells drive the formation of the HF and glial scaffold in the developing hippocampus, and *Tbr2* has been implicated in the regulation of migration of CR cells (Frotscher et al., 2003; Hodge et al., 2013; Meyer et al., 2004; Meyer et al., 2019). While in the CH, *Tbr2* is specifically expressed in differentiating CR cells, this gene is also expressed in hippocampal progenitors (that do not arise from the CH) (Hodge et al., 2013; Hodge et al., 2012). Interestingly, in *Lmx1a* mutants, *Tbr2* expression was reduced in the CH, but not in hippocampal progenitors (Fig. 4B-D). To study whether *Tbr2* regulates CR cell migration cell-autonomously, we conditionally inactivated this gene in the CH (in *Lmx1a-Cre:2Tbr2^F/F^* mice), which resulted in CR cell migration abnormalities similar to those observed in *Lmx1a-/-* mice and *Lmx1a/Tbr2* double heterozygous (*Lmx1a+/-;1.mxla-Cre;Tbr2^+/F^*) mice. In addition to CR migration abnormalities, the latter two mouse strains had similar HF and glial scaffold formation abnormalities. Taken together, our data indicate that in addition to regulating hippocampal development via Wnt signaling, *Lmx1a* also regulates DG morphogenesis by promoting CR cell migration, at least partially, via *Tbr2*.

Initiation of CR cell migration coincides with their differentiation in the CH (Causeret et al., 2021). In addition to CR cell migration, neuronal differentiation was also disrupted in the *Lmx1a* mutant CH, as revealed by our RNAseq pathway analysis and a delayed exit of CH progenitors from the cell cycle, a prerequisite for cell differentiation (Fig. 2). Interestingly, despite a delayed migration of CR cells, exit of progenitors from the cell cycle was normal in the CH of *Lmx1a+/-*; *Tbr2* CH null/+ double heterozygous mice (Fig. 4G-H), indicating that CR cell differentiation and migration are regulated by *Lmx1a* via different downstream mediators. We found that loss or knockdown of *Lmx1a* in the CH reduces the expression of a negative regulator of the cell cycle *Cdkn1a* (Xiao et al., 2020) and that exogenous *Cdkn1a*, introduced via *in utero* electroporation, not only rescued cell-cycle exit defects in the CH but also normalized the number of p73+ CR cells in *Lmx1a-/-* embryos. Together, our data indicate that *Lmx1a* promotes differentiation of CR cells at least partially via regulating the exit of their progenitors from the cell cycle via *Cdkn1a*.

Our current analysis has also strengthened the role of *Lmx1a* in the segregation of the CH from the cortical neuroepithelium. A boundary formation between the cortical (hippocampal) and CH neuroepithelia involves the cortical selector gene *Lhx2*. In the chimeric telencephalic neuroepithelium and *in vitro* cell aggregation studies, *Lhx2*+/+ and *Lhx2*-/- cells self-segregate, forming a boundary between *Lhx2*+/+ (*Lhx2* expressing) cells that adopt the hippocampal fate and *Lhx2-/-* cells that adopt the fate of the CH (Mangale et al., 2008). Our previous genetic fate mapping revealed that in the absence of *Lmx1a*, the dorsal midline lineage aberrantly contributes to the adjacent hippocampus (Chizhikov et al., 2010). Our current observation that *Lmx1a* is sufficient to downregulate *Lhx2* in e10.75-e12.5 cortical neuroepithelium (Fig. 6J-L) indicates an intrinsic role for *Lmx1a* in the segregation of the CH lineage from cortical (hippocampal) cells. Interestingly, our RNAseq analysis revealed significant enrichment in cell-cell and cell-extracellular matrix adhesion pathways among genes misregulated in the *Lmx1a-/-* CH, suggesting that in addition to *Lhx2*, *Lmx1a* regulates CH/hippocampus segregation via adhesion genes.

In conclusion, we determined that *Lmx1a* orchestrates CH development and function by co-regulating multiple processes ranging from CH cell-fate induction to CH Wnt signaling to differentiation and migration of CR cells. We determined the genes misregulated by loss of *Lmx1a*, and, by combining gene expression, genetic analysis and *in utero* electroporation rescue experiments, identified *Wnt3a, Tbr2*, and *Cdkn1a* as key downstream mediators of *Lmx1a* function in the CH. It is likely that genes identified as misexpressed in *Lmx1a-/-* CH by our RNAseq analysis include additional important regulators of CH development or mediators of CH function, and further studies are required to fully characterize an *Lmx1a*-dependent network. Regardless, our work revealed that the development and function of the CH, a key signaling center in the mammalian brain, employs *Lmx1a* as a master regulatory gene and established the framework for characterizing the mechanisms that regulate the development and function of signaling centers in the CNS.

## Materials and Methods

### Mice

Mice used in this study include *Lmx1a* null (*Lmx1a^drJ^*, Jackson Laboratory strain #000636) (Chizhikov et al., 2006; Deng et al., 2011; Kridsada et al., 2018), *Wnt3a* null (Jackson Laboratory strain #004581) (Takada et al., 1994), *Tbr2^floxed^* (Jackson Laboratory strain #017293) (Zhu et al., 2010), and *Lmx1a-Cre* (BAC transgenic mice that express a GFP-tagged Cre protein under the control of the *Lmx1a* regulatory elements, referred to as *Lmx1a-Cre* throughout the paper) (Chizhikov et al., 2010). Animals were maintained on a mixed genetic background comprising C57Bl6, FVB, and CD1. Genomic DNA obtained from mouse tails was used for genotyping (Chizhikov et al., 2019; Takada et al., 1994; Zhu et al., 2010). Both males and females were used for analysis. The presence of a vaginal plug was considered as embryonic day 0.5 (e0.5), while the day of birth was considered postnatal day 0 (P0). For cell proliferation, cell cycle exit, and cell migration experiments, pregnant dams were intraperitoneally injected with BrdU at 50mg/kg. All animal experiments were approved by the University of Tennessee Health Science Center (UTHSC) Institutional Animal Care and Use Committee (IACUC).

### Immunohistochemistry

Embryonic heads or dissected brains were fixed in 4% paraformaldehyde (PFA) in phosphate buffered saline (PBS) for 1.5 to 12 h, washed three times in PBS (30 min each), equilibrated in 30% sucrose in PBS for 2 h at 4°C, and embedded in optimum cutting temperature (OCT) compound (Tissue Tek). Tissue blocks were sectioned coronally on a cryostat at 12 μm, and sections were collected on Superfrost Plus slides (Fisher Scientific). Immunohistochemistry was performed as described (Iskusnykh et al., 2021). Sections were dried for 20 min, washed three times in PBS (10 min each), blocked in PBS containing 1% normal goat serum and 0.1% Triton X-100, and incubated with primary antibodies at 4°C overnight. Slides were washed three times in PBS (10 min each), incubated with secondary antibodies for 1 h at room temperature, washed three times in PBS (10 min each), mounted in Fluorogel (EMS), and cover slips were applied. For Ki67 and BrdU immunostaining, slides were boiled in 1x target retrieval solution (Dako) for 10 min and cooled at room temperature for 1 h prior to the blocking step. We used the following primary antibodies: anti-Prox1 (rabbit, Chemicon, catalog #Ab5475, 1:500, RRID:AB_177485), anti-BrdU (rat, Abcam, catalog #ab6326, 1:50, RRID:AB_305426), anti-BrdU (rabbit, Rockland, catalog #600-401-c29, 1:300, RRID:AB_10893609), anti-Ki67 (mouse, BD Pharmingen, catalog #556003, 1:250, RRID:AB_396287), anti-GFAP (rabbit, Dako, catalog #z0334, 1:200, RRID:AB_10013382), anti-Reelin (mouse, Chemicon, catalog #Mab5364, 1:500, RRID:AB_2179313), anti-Tbr2 (rat, eBioscience, catalog #14-487582, 1:200, RRID:AB_11042577), anti-GFP (chicken, Abcam, catalog #ab13970, 1:500, RRID:AB_300798, anti-p73 (mouse, Invitrogen, catalog #MA5-14117, 1:100, RRID:AB_10987160), anti-Lhx2 (rabbit, Invitrogen, catalog #PA5-64870, 1:200, RRID:AB_2662923), anti-Cdkn1a (mouse, BD Pharmingen, catalog #556431, 1:500, RRID:AB_396415), anti-Ctip2 (rat, Abcam, catalog #ab18465, 1:500, RRID:AB_2064130), anti-Lmx1a (goat, Santa-Cruz, catalog #sc-54273, 1:500, RRID:AB_2136830), and anti-activated Caspase 3 (rabbit, Promega, catalog #G7481, 1:250, RRID:AB_430875), and appropriate secondary antibodies, conjugated with Alexa 488 or 594 fluorophores (Life Technologies) at 1:100 dilutions.

### Histology

Tissue was processed as described for immunohistochemistry above. Slides were incubated in 0.1% cresyl violet acetate solution for 15 min, dehydrated in ethanol and xylenes, dried at room temperature, mounted in Permount (Fisher), and cover slips were applied.

### RNAscope fluorescent *in situ* hybridization

Embryos were fixed in 4% PFA for 16 h at 4°C, submerged in 10%, 20%, and 30% sucrose, embedded in OCT, and sectioned coronally on a cryostat at 12 μm. *In situ* hybridization was performed using an RNA probe for the mouse *Wnt3a* gene with 20 ZZ pairs (NM_009522.2, target location: 667 – 1634) and the RNAscope^®^ 2.5 HD Reagent Kit-RED (Advanced Cell Diagnostics, USA). Slides were boiled in 1X Target retrieval solution for 5 min, treated with protease for 15 min at 40°C, and incubated with the RNA probe for 2 h at 40°C. The unbound probe was removed by rinsing slides in 1X wash buffer, and signal amplification reagents were sequentially added. Finally, slides were washed twice in PBS, mounted in Fluoro-Gel medium, and cover slips were applied (Electron Microscopy Sciences, USA). When slides from *in utero* electroporated embryos were used, GFP fluorescence in each section was imaged to identify electroporated cells prior to the beginning of *in situ* hybridization.

### Laser capture microdissection (LCM) and RNA purification

LCM was performed using an Arcturus XT laser capture microdissection machine. For RNAseq experiments (Fig. 2G, H), the CH was isolated under a bright light mode. For *Lmx1a* knockdown experiments, achieved via *in utero* electroporation, electroporated (GFP+) cells in the CH were isolated using a fluorescent mode (see experimental design in Suppl. Fig. 5A-D).

For both types of experiments, embryonic heads were placed in OCT and flash-frozen on dry ice. Serial 10 μm thick coronal sections spanning the entire telencephalon were collected. Every second section was immunostained against Lmx1a to confirm the presence of the CH and identify its boundaries. The remaining slides were used to laser capture microdissect the entire CH (for RNAseq experiments) or GFP+ (electroporated) cells in the CH (for *Lmx1a* shRNA knockdown *in utero* electroporation experiments). CH tissue isolated from different sections of the same embryo was pooled and comprised one sample for RNAseq or qRT-PCR analysis. In *Lmx1a* knockdown experiments aimed to confirm an intrinsic role of *Lmx1a* in the CH, only embryos showing GFP fluorescence in the CH but lacking that in the ChPe (where Lmx1a is expressed in addition to CH (Chizhikov et al., 2019) were used for analysis (Suppl. Fig. 5B).

A Pico Pure RNA purification kit (Arcturus) was utilized to isolate RNA from laser capture microdissected samples, according to the manufacturer’s instructions. RNAse-free DNAse I (Qiagen) was used to remove traces of genomic DNA.

### RNAseq and bioinformatics analysis

All RNA samples were subject to quality control using a Bioanalyzer. Mean RNA integrity number (RIN) of the samples used for library construction and sequencing was 8.43+0.69 (sd). FastQC-v0.11.8 was used to perform quality control of the RNA-Seq reads. Remaining adapters and low-quality bases were trimmed by Trimmomatic-v 0.36 (Bolger et al., 2014) with parameters: Adapters:2:30:10 SLIDINGWINDOW:4:15 MINLEN:30. Paired-end RNA-sequencing data was aligned to the mouse genome (GRCm38) with gene annotations from Ensembl (v102) (Aken et al., 2016) using STAR-v2.7.1a with default parameters (Dobin et al., 2013). Read count per gene was calculated using HTSeq-v0.11.2 (Anders et al., 2015). Genes were filtered by requiring at least 10 read counts in at least three samples and then processed using EdgeR-v3.30.3 (Robinson et al., 2010) for normalized read count per million (CPM) and differential expression analysis. P-values were adjusted by the false discovery rate (FDR). Significant differentially expressed genes (DEGs) were declared at FDR < 0.1. Functional enrichment was done using R package clusterProfiler-v 3.18 (Yu et al., 2012) based on the biological processes Gene Ontology (GO), hallmark, and curated gene set definitions from MSigDB (v7.4.1) (Subramanian et al., 2005).

### qRT-PCR analysis

cDNA was synthesized with the cDNA synthesis kit (Bio-Rad, cat#1708890). qRT-PCR was performed using a Roche LC480 Real-time PCR machine and SYBR Fast qPCR master mix (Kapa Biosystems) as described (Iskusnykh et al., 2021). Gene expression was normalized to that of the reference gene *Gapdh*. All samples were tested in triplicate, using the primers listed below.

### Mouse *in utero* electroporation

For overexpression studies, cDNAs encoding full-length mouse *Lmx1a* and *Cdkn1a* were cloned into a *pCIG* plasmid (Megason and McMahon, 2002). For *Lmx1a* knockdown experiments, we used the previously characterized *Lmx1a*-targeting shRNA construct (*pLKO. 1*-mouse *Lmx1a* shRNA, Sigma, TRCN0000433282) (Fregoso et al., 2019). The *pLKO*. 1-non-mammalian shRNA construct (Sigma, SHC002) was used as a control. Plasmids were co-electroporated with *pCAG-GFP* (80) to visualize electroporated cells.

Using a fine glass microcapillary, plasmid DNA (1μg/μl for each plasmid) was injected into the lateral ventricles of the embryos. Electroporations were performed using paddle electrodes and a BTX ECM830 electroporator. The positive electrode was placed against the medial telencephalic neuroepithelium to target the CH or neighboring hippocampal primordium. Following electroporation, the uterus was placed back into the abdomen to allow continuing development. Upon harvesting, embryos were evaluated under a stereomicroscope for GFP fluorescence in the telencephalon. The telencephalon of GFP+ embryos was serially coronally sectioned, and those demonstrating GFP fluorescence in desired telencephalic areas were used for analysis.

### Electrophysiological Recordings

Brains were dissected in artificial cerebrospinal fluid (aCSF), which contained 20 mM D-glucose, 0.45 mM ascorbic acid, 1 mM MgSO_4_, 2.5 mM KCl, 26 mM NaHCO_3_, 1.25 mM NaH_2_PO_4_, 125 mM NaCl, and 2 mM CaCl_2_, and were cut into 300 μm thick slices using a Leica VT1000S vibratome at 4°C. Slices were allowed to recover at 32°C for 60 min in an aCSF-filled chamber under continuous bubbling with 95% O_2_/5% CO_2_. For the recording, slices were put in a chamber attached to a modified stage of an Olympus BX51WI upright microscope and perfused continuously with bubbled aCSF at room temperature. Electrodes (4–8 MΩ) were pulled from borosilicate glass with an outer diameter of 1.5 mm using a P-97 micropipette puller (Sutter Instruments, USA) and filled with a solution containing 140 mM K-gluconic acid, 10 mM HEPES, 1 mM CaCl_2_, 10 mM EGTA, 2 mM MgCl_2_, and 4 mM Na2ATP (pH 7.2, 295 mOsm). Whole-cell current clamp recordings were obtained from neurons in the granule cell layer of the DG using an Axon Multiclamp 700B amplifier (Molecular Devices, USA). Traces were digitized using an Axon 1440A Digitizer at 10 kHz and filtered at 2 kHz using Clampex 10 software (Molecular Devices, USA). After achieving a stable whole-cell recording, a multi-sweep current injection step protocol (−120 to 20 pA in 20 pA increments) was applied, and input resistance was calculated using Clampfit 9 software (Molecular Devices, USA).

### Measurements, cell counts, and statistical analysis

Serial coronal sections were generated for each embryonic or postnatal telencephalon. The same number of sections was collected on each slide to facilitate the identification of the position of each section along the anterior-posterior (A-P) axis of the telencephalon. Upon inspection under a bright field microscope, sections containing the CH (or its derivative – the fimbria in late embryonic and postnatal brains) were identified. For consistency, we compared sections from control and experimental animals taken at similar A-P levels, approximately midway along the telencephalic region that contained the CH/fimbria. At least three embryos were analyzed per experimental condition (typically three-five non-adjacent sections per animal).

Sections containing a fluorescent signal were evaluated under a Zeiss Axio Imager A2 fluorescence microscope and imaged using an AxioCam Mrm camera and AxioVision Rel 4.9 software (Zeiss). Bright-field images were taken using an Olympus SZX microscope, Leica MC170HD camera and LAS V4.5 software. In all experiments, slides for control and experimental animals were processed and stained in parallel and imaged using identical acquisition settings. Signal intensity (average pixel intensity) was measured in ImageJ software (NIH), as previously described (Chizhikov et al., 2019). Wnt3a RNAscope *in situ* hybridization signal was quantified in the entire CH (Fig. 3C). Immunohistochemistry signal for Tbr2 and Cdkn1a was quantified in all positive cells located in the CH (Figs. 4C, 5D) or in a 200 μm high segment of the hippocampal primordium located directly above the CH (Fig. 4D). Immunohistochemistry signal for Lhx2 was measured in all GFP+ cells in a 200 μm high segment of the hippocampal primordium located directly above the CH (Fig. 6L). The signal intensity in individual cells was averaged to generate a value for each embryo. For these and other experiments, the boundaries of the CH were identified using adjacent Lmx1a-immunostained sections as templates (taking advantage that in the *dreher^J^* strain used in this study, *Lmx1a* harbors a point mutation, which although completely functionally inactivates Lmx1a (Chizhikov et al., 2010; Deng et al., 2011) does not prevent the production of (inactive) Lmx1a protein).

To analyze proliferation, a 1.5-hr BrdU pulse (which labels cells in the S phase of the cell cycle) was used, and the percentage of BrdU+ cells (the number of BrdU+ cells divided by the total number of cells in the region of interest) was calculated. The percentage of cells that exited the cell cycle (the number of BrdU+/Ki67-cells divided by the total number of BrdU+ cells in the region of interest) was calculated in embryos 24 h after BrdU injection as described (30). For migration experiments, mice were injected with BrdU at e14.5, shortly before the DNe progenitors begin migration to the DG, and the embryos were collected at e18.5. The fractions of BrdU+ cells located in three segments along their migratory route were calculated (Fig. 1K-M): the first segment (labeled as DMS) was between the DNe and FDJ, the second was along the surface of the FDJ, and the third was in the DG (Cai et al., 2018).

Comparisons of two groups were performed with an unpaired two-tailed t-test. Comparisons of multiple groups were performed with one-way ANOVA with Tukey’s post hoc test. IBM SPSS Statistics (version 27) and Excel software were used. Quantitative data in all figures, except Figures 4A, 5A, and Suppl. Figures 4C and 6B, which show RNAseq expression data in a log scale, are presented as the mean ± standard deviation (sd). In the aforementioned figures, data are presented as the median (center line), the first and third quartiles (upper and lower edges of the box, respectively), and error bars indicating the minimum and maximum data points. Graphs were generated with GraphPad Prism 6 (Graphpad) software.

## Acknowledgments

We thank W. Armstrong, M. Ennis, R. Foehring, T. Ishrat, D. Guan, and L. Wang for advice on experimental protocols or sharing equipment, S. Krat, L. Mukhametzyanova, and A. Zakharova (UTHSC) for genotyping, and K. Johnson-Moore and the UTHSC Office of Scientific Writing for manuscript editing. This work was supported by NIH R01 NS093009 to V.V.C and UTHSC Neuroscience Institute.

## Author Contributions

I.Y.I., N.F., and V.V.C. designed the study and wrote the paper; I.Y.I, N.F., Y.L., L.B., M.K.K, E.Y.S., P.A.N., and V.V.C. performed experiments and/or analyzed data.

