## Supplemental information for "Lmx1a is a master regulator of the cortical hem"

Iskusnykh et al.

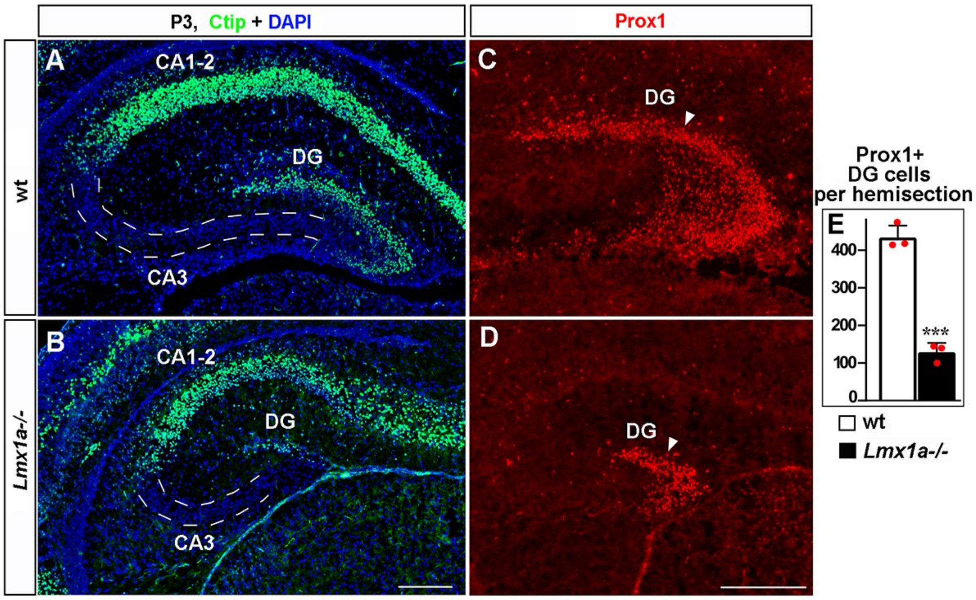


**Suppl. Fig. 1. Normal hippocampal patterning but a reduced number of DG neurons in *Lmx1a-/-* mice at P3.**
(A, B) Ctip2 immunohistochemistry revealed normal relative location of the DG, CA3 and CA1-2 hippocampal regions in *Lmx1a-/-* mice.
(C-E) The number of Prox1+ DG neurons (arrowhead) was reduced in P3 *Lmx1a-/-* mice. ***p<0.01, n=3 mice per genotype.
Scale bars: 200 µm.

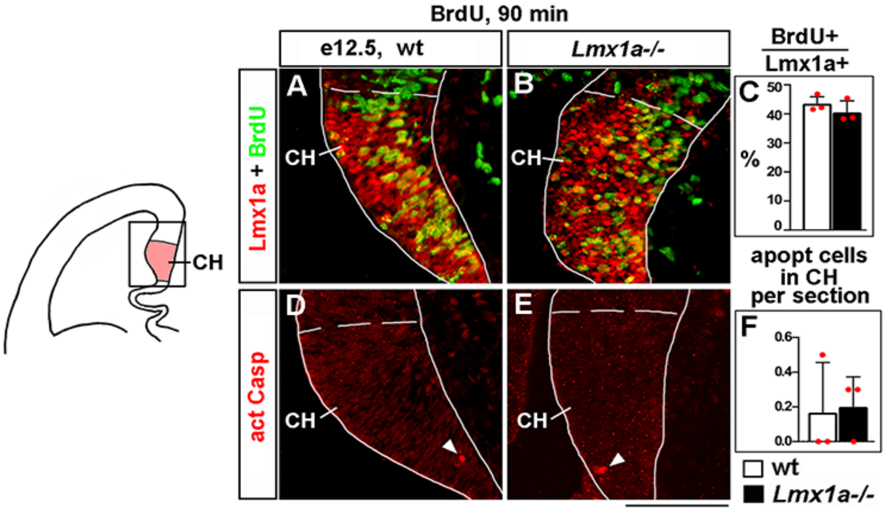


**Suppl. Fig. 2. Normal proliferation and apoptosis in the CH of *Lmx1a-/-* embryos at e12.5.**
(A-C) Proliferation (% of BrdU+ cells in the CH after a 90 min BrdU pulse) was similar in wild type and *Lmx1a-/-* embryos. Lmx1a immunohistochemistry was used to identify the CH (outlined by dashed line). (Note that in our *Lmx1a-/-* mice, *Lmx1a* is inactivated by a point mutation ($dr^J$ allele) (Chizhikov

et al., 2006; Deng et al., 2011), which leads to the production of inactive Lmx1a protein, detectable by immunohistochemistry).
(D-F) The number of apoptotic (activated Caspase+) cells in the CH was similar in *Lmx1a-/-* and wild type embryos. n=3 embryos per genotype.
Scale bars: 100 µm.

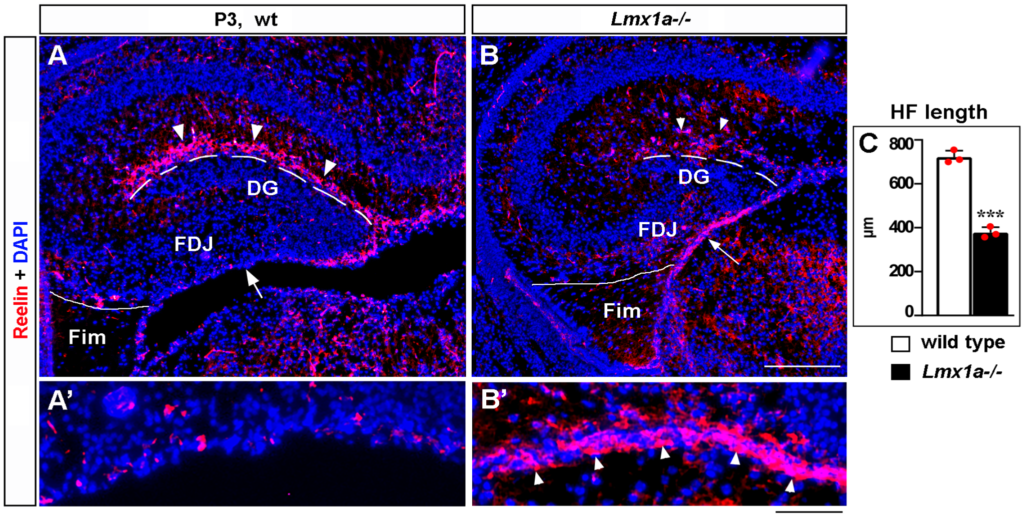


**Suppl. Fig. 3. Abnormal distribution of CR cells in P3 *Lmx1a-/-* mice.**
(A-B') In P3 wild type mice, many CR cells populate the HF, and virtually none of them remain at the FDJ surface (A, A'). In contrast, many CR cells were still located at the FDJ surface in *Lmx1a-/-* littermates (B, B'), which was associated with a reduced HF length (dashed line in A, B; C). ***p<0.001, n=3 mice per genotype.
Fim – fimbria; FDJ – fimbrial-dentate junction. FDJ surface is pointed by arrow in a, b and is shown at higher magnification in A', B'. Arrowheads point to (Reelin+) CR cells.
Scale bars: 200 µm (A, B); 50 µm (A', B').

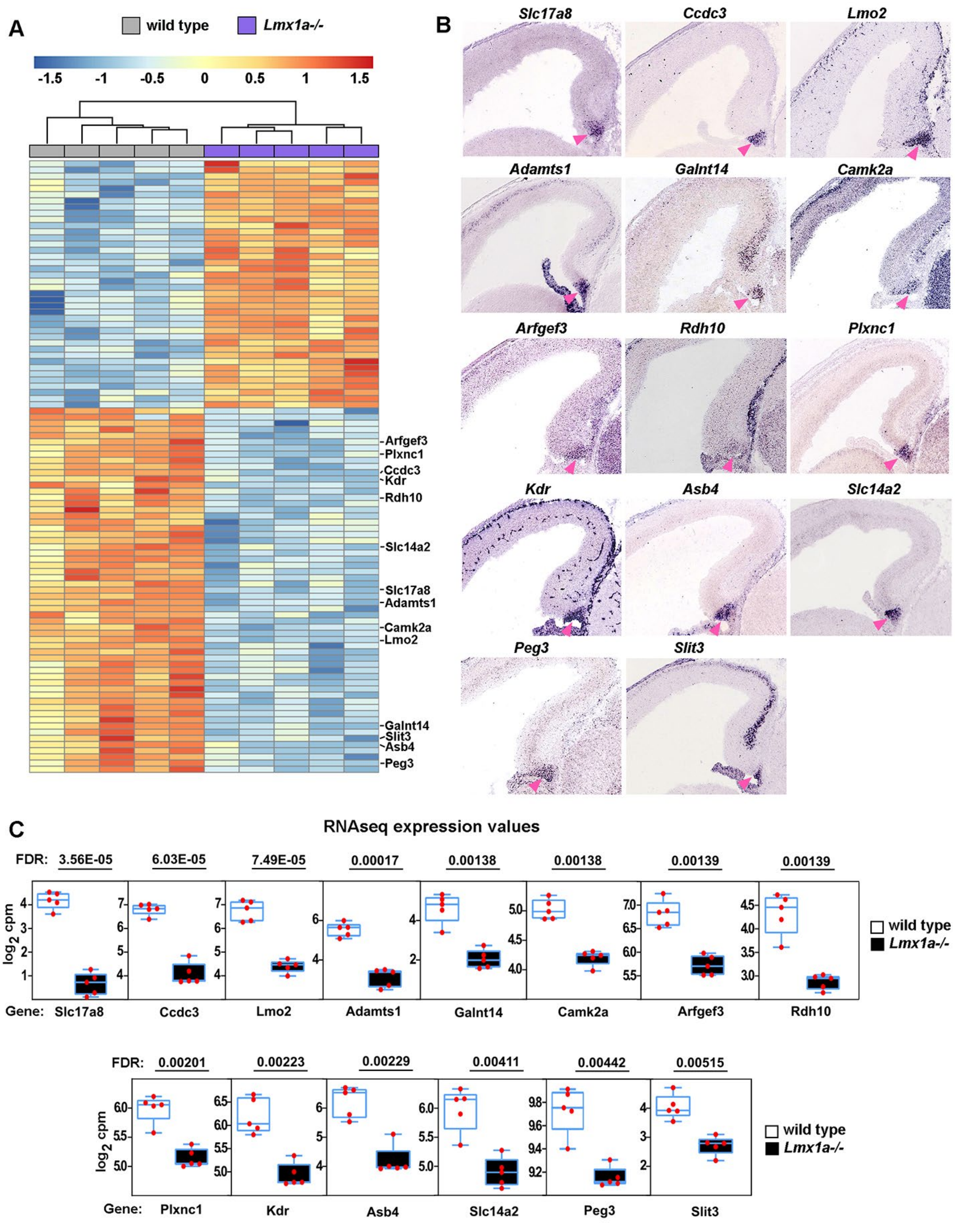
A
wild type
Lmx1a-/-
-1.5 -1 -0.5 0 0.5 1 1.5
Arfgef3
Plxnc1
Ccdc3
Kdr
Rdh10
Slc14a2
Slc17a8
Adamts1
Camk2a
Lmo2
Galnt14
Slit3
Asb4
Peg3
B
Slc17a8
Ccdc3
Lmo2
Adamts1
Galnt14
Camk2a
Arfgef3
Rdh10
Plxnc1
Kdr
Asb4
Slc14a2
Peg3
Slit3
C
RNAseq expression values
FDR: 3.56E-05 6.03E-05 7.49E-05 0.00017 0.00138 0.00138 0.00139 0.00139
log2 cpm
Gene: Slc17a8 Ccdc3 Lmo2 Adamts1 Galnt14 Camk2a Arfgef3 Rdh10
wild type
Lmx1a-/-
FDR: 0.00201 0.00223 0.00229 0.00411 0.00442 0.00515
log2 cpm
Gene: Plxnc1 Kdr Asb4 Slc14a2 Peg3 Slit3
wild type
Lmx1a-/-

**Suppl. Fig. 4. Reduced expression of CH markers in *Lmx1a-/-* embryos.**
(A) Hierarchical clustering heatmap of 100 transcripts whose expression is most significantly misregulated in the CH of e13.5 *Lmx1a-/-* embryos relative to wild type littermates. A search of publically accessible *in situ* hybridization databases identified 14 of these transcripts as specifically or predominantly expressed in the CH (listed on the right).
(B) *In situ* hybridization images showing predominant CH expression of these 14 genes in e14.5 wild type embryos. The CH is indicated by the pink arrowhead. All the images are from the Gene Paint database (Visel et al., 2004; https://gp3.mpg.de/).
(C) Normalized read counts from the RNAseq experiment (Fig. 2G, H) for the above genes. Genes are listed from most to least significantly misregulated in the CH of *Lmx1a-/-* embryos. FDR values are shown above the genes.

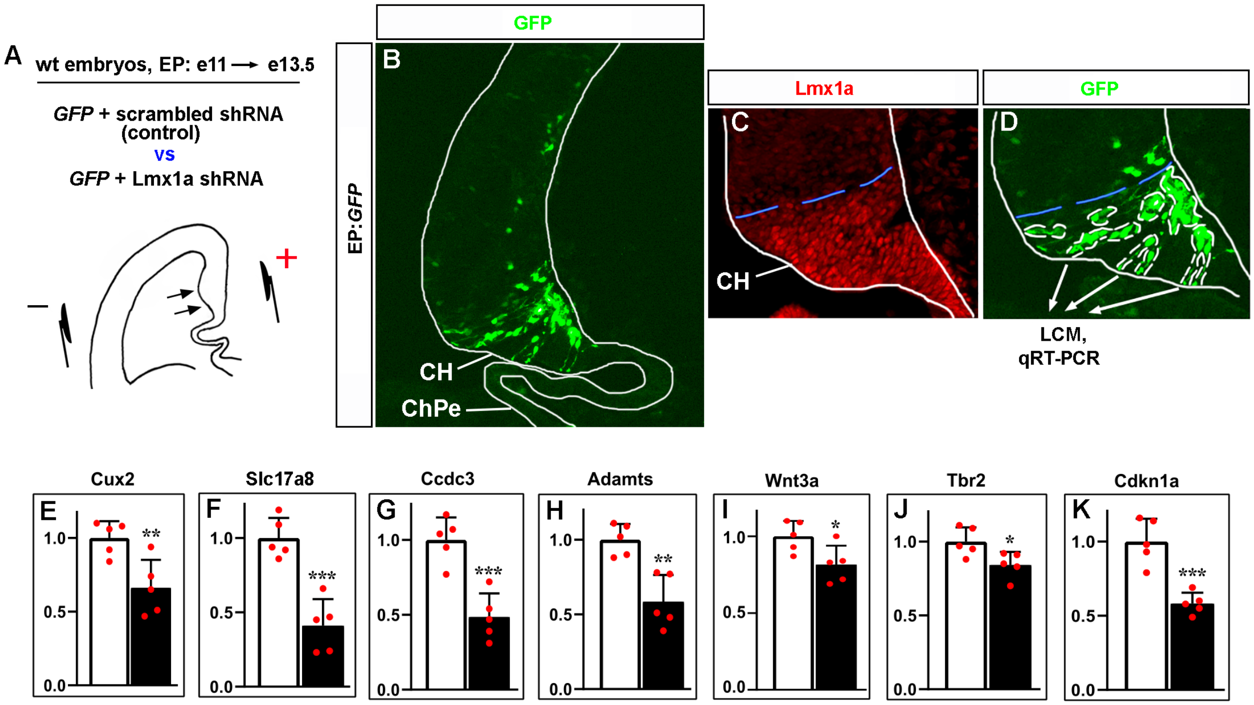


**Suppl. Fig. 5. *Lmx1a* downregulation specifically in the CH reduces the expression of CH markers and key CH developmental genes.**
(A-D) Experimental design. The CH area of wild type e11 embryos was *in utero* electroporated with plasmids encoding GFP+scrambled shRNA (controls) or GFP + previously validated Lmx1a shRNA (4) (experimental embryos), by placing electrodes as shown in the schematic (A). Negatively charged DNA moves toward the positive electrode (arrows) entering the CH. Embryos were collected at e13.5, serially coronally sectioned, and those demonstrating GFP expression in the CH but not in the ChPe (the only telencephalic structure beyond the CH that expresses *Lmx1a*) were used for analysis (see an example in panel B). All GFP+ cells in the CH (outlined by white dotted lines in panel D) from five sections per embryo were isolated by LCM and used for qRT-PCR analysis. The blue dashed line shows the dorsal boundary of the CH, which was determined using adjacent Lmx1a-immunostained sections (C). Panel D shows a higher magnification of the CH area of the section shown in panel B.

(E-K) qRT-PCR analysis revealed that targeting of *Lmx1a* in the CH reduces expression of key CH markers and CH developmental genes. p values are shown above each gene. ***p<0.001, **p<0.01, *p<0.05. n=5 embryos per condition. A direct Lmx1a downstream target *Cux2* (Fregoso et al., 2019) was used as a positive control to confirm the disruption of Lmx1a function in these experiments (E).

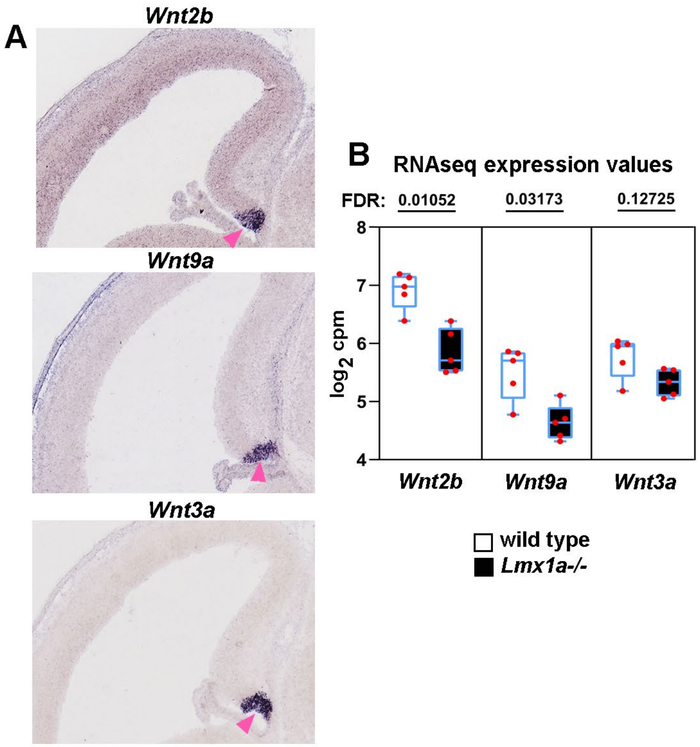


**Suppl. Fig. 6. Canonical Wnts misregulated in the CH of *Lmx1a-/-* embryos based on RNAseq analysis.**
(A) *In situ* hybridization images showing that in the telencephalon, *Wnt2b*, *Wnt9a*, and *Wnt3a* are specifically expressed in the CH (pink arrowhead). All images show wild type e14.5 embryos from the Gene Paint database (Visel et al., 2004; https://gp3.mpg.de/).
(B) Normalized read counts from the RNAseq experiment (Fig. 2G, H) for the above Wnts. FDR values are shown above the genes.

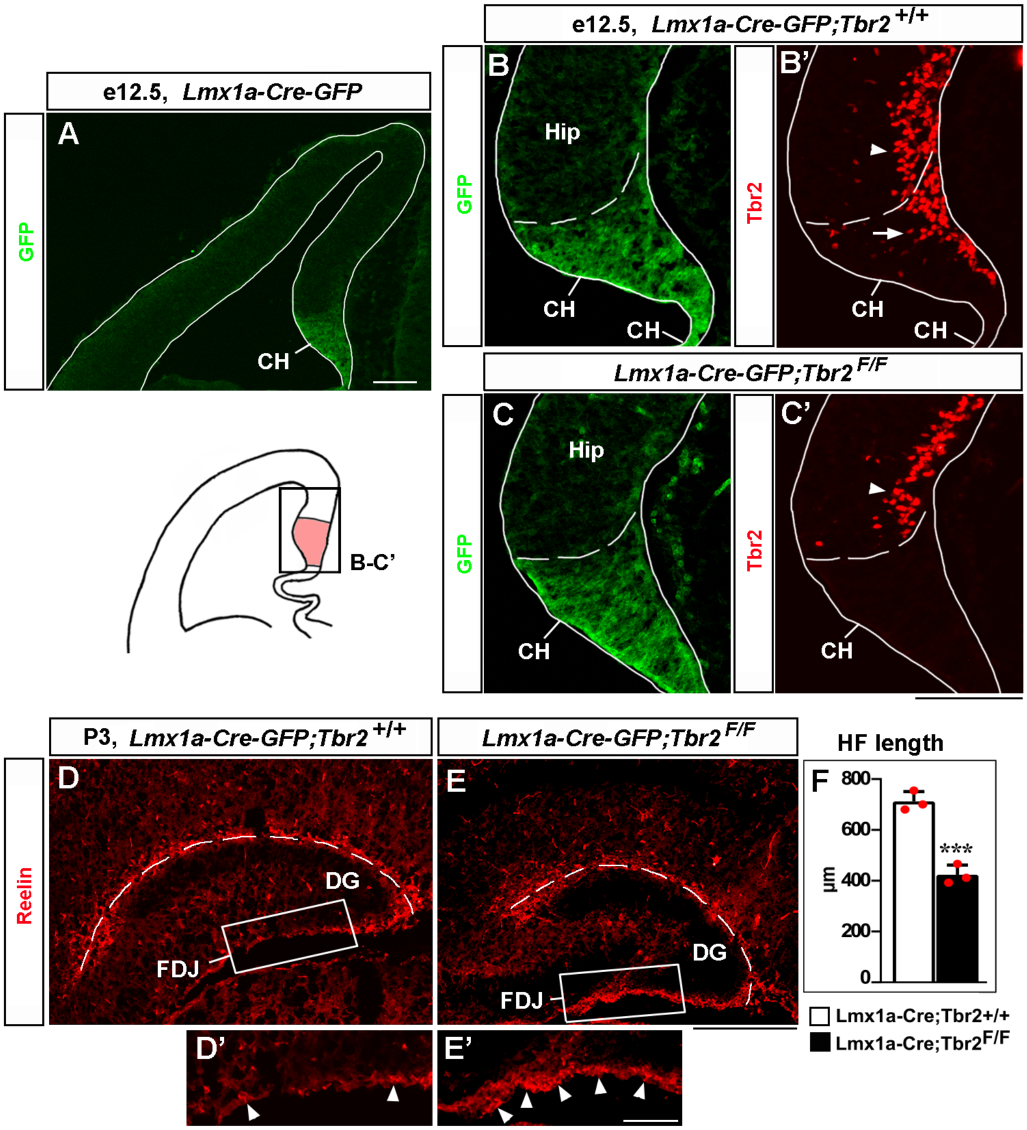


**Suppl. Fig. 7. Loss of *Tbr2* in CH-derived CR cells compromises their migration and HF formation.**

(A) In the telencephalic neuroepithelium of *Lmx1a-Cre-GFP* mice, *Cre-GFP* expression is limited to the CH (Chizhikov et al., 2010).

(B-C') Specific deletion of *Tbr2* in the CH. Panels B and B' and panels C and C' show the same sections imaged for different antibodies. Dashed line demarcates the CH from the hippocampal primordium (Hip). Arrow points to Tbr2 expression in CR cells in the CH (Hodge et al., 2013) of control (*Lmx1a-Cre-GFP;Tbr2*$^{+/+}$) embryos (B'). Tbr2 expression is lost in the CH of *Lmx1a-Cre-GFP;Tbr2*$^{F/F}$ embryos (C'). *Tbr2* expression is still present in hippocampal intermediate progenitors (Hodge et al., 2013) in both control and *Lmx1a-Cre-GFP;Tbr2*$^{F/F}$ embryos (arrowhead in B', C').

(D-F) In P3 *Lmx1a-Cre-GFP;Tbr2*$^{F/F}$ mice, excessive Reelin+ CR cells accumulate at the FDJ surface (D', E', arrowheads), which was associated with a reduced HF length (dashed line in D, E; F). ***$p<0.001$.

Scale bars: 100 µm (A-C'); 200 µm (D, E); 50 µm (D', E').

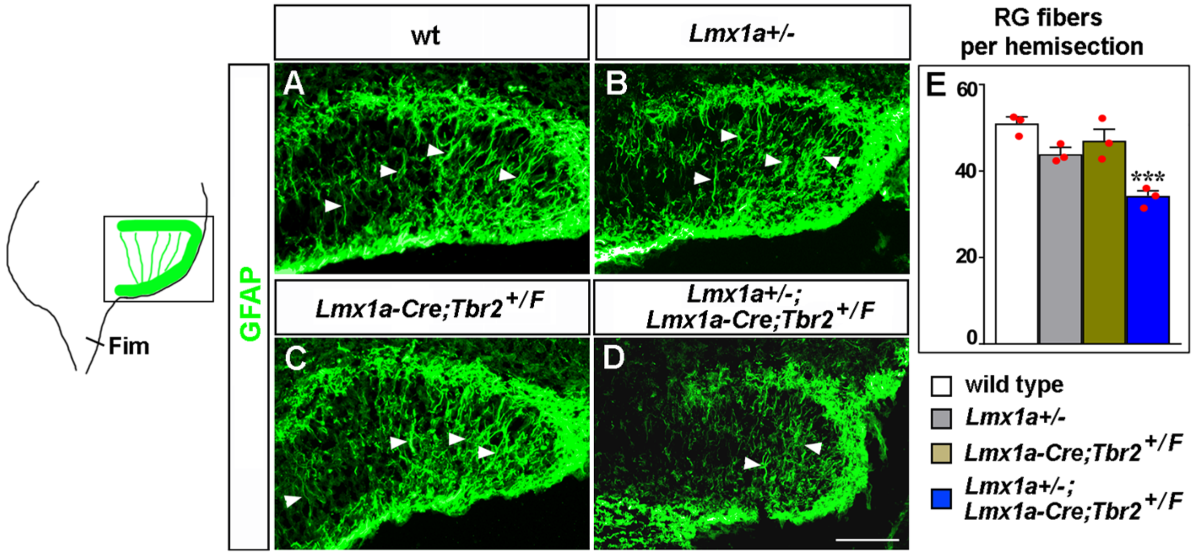


**Suppl. Fig. 8. *Lmx1a* regulates the transhilar glial scaffold via *Tbr2*.**
(A-E) The number of GFAP+ transhilar glial fibers was reduced in *Lmx1a/Tbr2* double heterozygotes (*Lmx1a+/-;Lmx1a-Cre;Tbr2*$^{+/F}$ mice – *Lmx1a+/-* mice in which one copy of *Tbr2* was deleted specifically in CH), but not single-gene heterozygotes, compared to wild type controls at e18.5. ***p<0.001, n=3 mice per genotype. Arrowheads indicate GFAP+ glial fibers that cross the hilus. Scale bars: 100 µm.

**Suppl. Table 1. Sequences of primer used in qRT-PCR**

| Primers | Reference |
|---|---|
| Slc17a8 F: ggtgtggggaccctctctgg<br><br>Slc17a8 R: cccagaagcgaagaccccgt | This study |
| Ccdc3 F: tatgccaaggtgctggcgct<br><br>Ccdc3 R: taaggttgagccgggagccg | This study |
| Adamts1 F: ctggcacctccggtggctta<br><br>Adamts1 R: gtccccatggtccccagctt | This study |
| Tbr2 F: ctacgggccatacgccggaa<br><br>Tbr2 R: gtagtgggcggtggggttga | This study |
| Cux2 F: ccctgaggaagacccctcgg<br><br>Cux2 R: ccttggcccatcaggaccca | This study |
| Cdkn1a F: ggtcccgtggacagtgagca<br><br>Cdkn1a R: gggacccagggctcaggtag | This study |
| Wnt3a F: ctcctctcggatacctcttagtg<br><br>Wnt3a R: gcatgatctccacgtagttcctg | Watanabe et al., 2016 |
| Gapdh F: cgacttcaacagcaactcccactcttcc<br><br>Gapdh R: tgggtggtccagggtttcttactcctt | Liu et al., 2009 |
